# MAPK Signaling and Angiopoietin-2 Contribute to Endothelial Permeability in Capillary Malformations

**DOI:** 10.1101/2025.03.31.646063

**Authors:** Sana Nasim, Mariam Baig, Jill Wylie-Sears, Matthew Vivero, Patrick Smits, Leanna Marrs, Yu Sheng Cheng, Cesar Alves, Anna Pinto, Arin K. Greene, Joyce Bischoff

## Abstract

Capillary malformations (CM) are slow-flow vascular abnormalities present at birth and predominantly manifest as cutaneous lesions. In the rare neurocutaneous disorder known as Sturge Weber Syndrome (SWS), individuals exhibit CM not only on the skin but also within the leptomeninges of the brain and the choroid of the eye. >90% of CM are caused by a somatic R183Q mutation in *GNAQ,* the gene encoding Gαq – a heterotrimeric G-protein subunit. The somatic *GNAQ* mutation is notably enriched in endothelial cells (ECs) isolated from CM-affected regions. Here we show blood vessels in cutaneous and leptomeningeal SWS lesions exhibit extravascular fibrin indicating a compromised endothelial barrier. Longitudinal MRI of the brain in one SWS patient further suggests vascular permeability. To explore this pathological phenotype, we employed the trans-endothelial electrical resistance (TEER) assay to measure permeability of the EC-EC barrier *in vitro*. Human EC CRISPR edited to create a *GNAQ* R183Q allele (EC-R183Q) exhibited a reduced barrier compared to mock edited EC (EC-WT). We sought to identify signaling molecules needed for EC barrier formation. Knockdown of angiopoietin-2 (ANGPT2), known to be significantly increased in EC-R183Q and in CM, partially yet significantly restored the barrier, while an anti-ANGPT2 function blocking antibody did not. We next tested the MEK1,2 inhibitor (Trametinib) because MAPK signaling is increased by *GNAQ* mutation. MEK1,2 inhibitors partially restored the EC barrier, implicating involvement of MAPK/ERK signaling. The combination of ANGPT2 knockdown and Trametinib significantly restored the EC barrier to near EC-WT levels. The additive impacts of ANGPT knockdown and MEK1,2 inhibition indicate the two operate in separate pathways. In summary, we discovered that *GNAQ* p.R183Q ECs exhibit compromised endothelial barrier formation, reflecting the compromised EC barrier in CM lesions, and that ANGPT2 knockdown combined with Trametinib effectively restores the EC-EC barrier.

**NONSTANDARD ABBREVIATIONS AND ACRONYMS:** 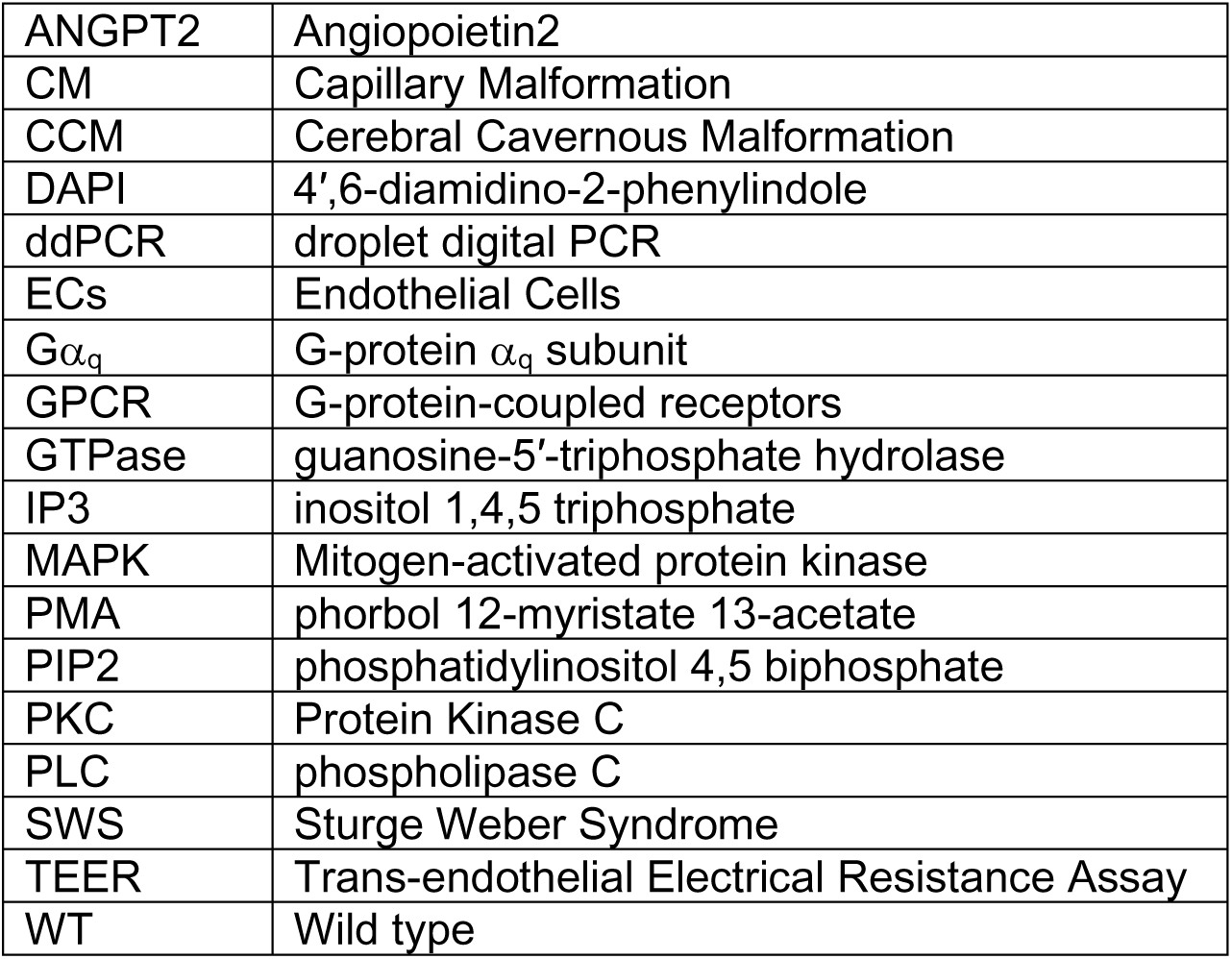

**NOVELTY AND SIGNIFICANCE:** *What is known?:*

1. The mutant Gαq-R183Q in endothelial cells activates phospholipase β3, contributing to increased angiopoietin-2, a pro-angiogenic, proinflammatory molecule that contributes to vascular permeability.
2. Endothelial Gαq-R183Q is sufficient to drive formation of enlarged blood vessels akin to what is observed in CM. ANGPT2 shRNA knockdown prevented the enlarged vessel phenotype in a xenograft model.
3. An EC-specific GNAQ p.R183Q mouse model showed permeability in brain vessels, detected by perfusion of Evans Blue dye, indicating reduced vascular integrity.

*What New Information Does This Article Contribute?:*

1. Reduced vascular integrity in CM is confirmed by Martius Scarlet Blue staining and longitudinal MRI imaging of SWS brain.
2. *GNAQ* p.R183Q EC form a weaker endothelial barrier *in vitro* compared to control ECs. The weakened endothelial barrier in the mutant ECscan be rescued by Gαq inhibitor, YM254890, confirming the compromised barrier is a consequence of the mutant Gαq.
3. Titration experiments modeling the mosaic nature of the *GNAQ* p.R183Q in CMshow that 5- 10% *GNAQ* p.R183Q EC in the monolayer is sufficient to reduce endothelial barrier formation.
4. Knockdown of ANGPT2 or MEK1,2 inhibition partially restored the endothelial barrier in *GNAQ* p.R183Q EC.
5. Combining knockdown of ANGPT2 and addition of a MEK inhibitor, Trametinib, restored the endothelial barrier to near what is seen in wild type ECs.

*What is the translational message?:* Sturge Weber Syndrome (SWS) is a neurocutaneous disorder that involves atypical blood vessel overgrowth in the skin, brain and eye. It is associated with facial CM (aka port wine birthmark), leptomeningeal CM in the brain visible with MRI, and glaucoma. Theneurological sequalae involve seizures, cerebral atrophies and calcification, and intellectual disorders. Currently there are no molecularly targeted therapies for non-syndromic CM or SWS. Our study shows the involvement of MAPK pathway and the proinflammatory molecule ANGPT2 in endothelial permeability and suggests a path to target *GNAQ* p.R183Q driven CM.

## INTRODUCTION

Capillary malformation (CM), also known as port-wine birthmark, is a slow flow vascular malformation consisting of abnormally enlarged blood vessels with uneven mural cell coverage^1-4^. In most cases, CM are non-syndromic and confined to skin. In the rare neurocutaneous disorder Sturge Weber Syndrome (SWS), facial CMs coincide with CM in the leptomeninges and the choroid of eye^5^. The majority of SWS patients are at high risk for seizures, stroke-like episodes, as well as cognitive impairment^6^. Only symptomatic treatments are available and to date there are no targeted molecular therapies for non-syndromic or syndromic CM.

A breakthrough study in 2013 discovered a somatic mutation in *GNAQ* (p.R183Q) in 92% of non-syndromic skin CMs and 88% of SWS skin and brain specimens^7^. The mutant allelic frequency, also referred to as variant allelic frequency, in CM affected tissues ranged from 1% to 18.1%^7^, underscoring the mosaicism of the somatic mutation. To date, multiple studies have confirmed the presence of the *GNAQ* p.R183Q mutation and shown variant allelic frequency between 1.2% to 33.3%^8-10^. A somatic missense mutation (p.R183C) in *GNA11*, a *GNAQ* homologue, was found in patients with diffuse CM on extremities, which have been reported to darken with time and exhibit milder neurological symptoms, suggesting a possible genotype- phenotype correlation^11,12^. Another somatic missense mutation in *GNB2* (p.K78E), the gene encoding the β subunit of the heterotrimeric G protein, was recently reported in a patient with SWS^13^. Our laboratory showed the *GNAQ* p.R183Q is enriched in endothelial cells (ECs) in skin and brain CMs^14,15^.

*GNAQ* encodes Gαq, an alpha subunit of the heterotrimeric G protein that links G-protein-coupled receptors (GPCR) to primarily activate phospholipase C (PLC)β3. GPCR activation causes Gαq to dissociate from the βγ subunits^16^, and increase the catalytic rate constant of PLCβ3^17^. Activated PLCβ3 hydrolyzes phosphatidylinositol 4,5 biphosphate (PIP2) to generate diacylglycerol (DAG) and inositol 1,4,5 triphosphate (IP3). PIP2 has long been established as a major contributor to the intracellular signaling stimulated by PLCβ^18^. DAG activates protein kinase C (PKC), which plays a role in various cellular processes including activation of downstream mitogen-activated protein kinase (MAPK) signaling^17^. IP3 acts as a second messenger, triggering the release of calcium ions from intracellular stores in the endoplasmic reticulum, which also leads to signaling and cellular responses. Gαq has intrinsic guanosine-5′- triphosphate hydrolase (GTPase) activity. Loss of arginine (R) at position 183, located in the conserved GTP-binding domain, is predicted to reduce GTPase activity and thereby increase GTP bound Gαq, which results in increased activity and increased signaling^19^.

The *GNAQ* p.Q209L mutation in uveal melanoma, an aggressive cancer in the eye, causes a stronger activation of the Gαq GTPase activity^20^. In uveal melanoma, ∼95% of the *GNAQ* or *GNA11* mutations are p.Q209L, while the p.R183Q mutation is found in 5% of uveal melanoma cases. Even though *GNAQ* p.R183Q is considered oncogenic in uveal melanoma, there is no clinical evidence of metastatic dissemination in CM. A protein-protein interaction network analysis suggests that p.R183Q and p.Q209L may have distinct downstream effectors leading to distinct cellular responses^21^. Nonetheless, the p.Q209L variant provides a template for understanding the consequences of these activating mutations^22^. Known downstream effecters of Gαq-Q209L are PLC/PKC, Rho/Rac signaling, and MAPK signaling including pERK, p38, Yap, and other signaling factors^22-25^. In contrast, the pathogenic downstream signaling pathways activated by *GNAQ* p.R183Q are not fully delineated. This gap in knowledge needs to be closed to advance our understanding of CM in both non-syndromic and syndromic SWS cases.

Previous studies using lentiviral constructs to express the p.R183Q mutation in normal human ECs showed activation of PLCβ3 and PKC, leading to increased angiopoietin-2 (ANGPT2), a potent angiogenic factor that promotes vascular permeability and remodeling. Furthermore, the p.R183Q ECs formed enlarged CM-like vessels in a murine xenograft model. Suppression of ANGPT2 using short hairpin RNA prevented the enlarged vessel phenotype, indicating its role in CM^19^. Increased ANGPT2 has been reported in other cellular models of *GNAQ* p.R183Q ECs as well^1,19^. Analysis of brain histopathology sections has shown that CM are associated with increased extravascular fibrin surrounding the enlarged and irregular blood vessels^4^. Moreover, a newly developed mouse model for GNAQ p.R183Q shows significant Evans Blue permeability in the brains of mutant mice^26^. To study the apparent increase in vascular permeability we analyzed the ability of p.R183Q expressing ECs to form electrically resistant endothelial barriers *in vitro*. Consistently, p.R183Q expressing ECs formed significantly weaker endothelial barriers, i.e., showed increased permeability, compared to wild-type (WT) ECs. Furthermore, we modeled the somatic mosaicism in CM by titrating EC-R183Q into EC-WT cultures and learned that a small percentage of mutant ECs have an outsized effect on the endothelial barrier. We used shRNA and siRNA to examine roles of ANGPT2 and MEK1 (aka MAP2K1) in endothelial barrier formation, and tested drugs against these molecules as orthogonal approaches. Taken together, our study pinpoints MEK1 and ANGPT2 as potential targets to reverse the cellular consequences of the mutant Gαq in CMs.

## METHODS

### Martius Scarlet Blue Staining

5µm sections of human brain and skin specimens were obtained from the Boston Children’s Hospital Department of Pathology under a human subject protocol (IRB-P00003505) approved by the Committee on Clinical Investigation at the Boston Children’s Hospital. Tissue sections were sent for Martius Scarlet Blue staining to iHisto (Salem, MA). MoticEasyScan Infinity 60 microscope was used for bright field scanning of each tissue section slide.

### MRI

Brain MRI was performed with 3-T Tim Trio MRI scanner (Siemens). Imaging protocol included sagittal volumetric T1-weighted images (magnetization-prepared rapid gradient echo [MPRAGE]), axial and coronal T2-turbo spin echo, axial fluid-attenuated inversion recovery, axial diffusion-weighted imaging, susceptibility-weighted imaging, and axial volumetric gadolinium-enhanced fat-suppressed T1-weighted images (MPRAGE). The retrospective study was approved by Boston Children’s Hospital IRB-P00014482.

### Droplet digital PCR

ddPCR was used to measure the mutant allelic frequency in tissue sections and in cultured cells as previously described^4,27^. Briefly, genomic DNA was isolated from FFPE tissue sections using a FormaPure kit (Beckman Coulter, C16675) or from cell pellets using DNeasy Blood & Tissue Kit (Qiagen, 69506). ddPCR with probes to detect the *GNAQ* R183Q mutation are listed in Major Resource Table. 15-30ng DNA per sample was mixed with ddPCR SuperMix (BioRad, 186-3010), primers for a 74bp amplicon in GNAQ Exon4, FAM-labeled quenched mutant probes and HEX labeled quenched wild-type probes (See Major Resource Table), divided into droplets (QX200 AutoDG) and amplified for 40 PCR cycles using 60°C as annealing temperature. The droplet fluorescence was read in a QX200 droplet reader and analyzed in QuantaSoft software (BioRad).

### Generation of GNAQ p.R183Q ECs

CRISPR-Cas9 gene editing was used to introduce the GNAQ p.R183Q mutation into htert-immortalized human aortic endothelial cells (teloHAECs, ATCC CRL-4052). Following IDT Alt-R Homology Directed Repair (HDR) protocol, TeloHAECs were nucleofected using Lonza 4D-Nucleofector (V4XP-5024, Lonza, Lexington, MA) with a solution containing preformed ribonucleoprotein (RNP) complexes combining IDT manufactured Alt-R sgRNA and Alt-R spCas9 enzyme (commercially available from Integrated DNA Technologies, Coralville, IA); homology directed repair (HDR) templates (IDT) for both GNAQ p.R183Q mutant and endogenous alleles and a GFPmax plasmid (available within Lonza V4XP- 5024 kit). HDR templates contained ∼50bp homology arms and three 3’- silent nucleotide changes just downstream to the *GNAQ* p.R183Q to prevent repeated Cas9 cleavage and increase recombination efficiency (see Major Resource table for full sequences). Nucleofection mixtures were supplemented with IDT electroporation enhancer buffer per IDT HDR Alt-R system protocol recommendations.

Following nucleofection, cells were grown at >20k cell/cm^2^ density overnight in Vasculife VEGF complete growth media (LL-0003, Lifeline Technologies, Frederick, MD) supplemented with 10% FBS (Gibco, Life Technologies, Carlsbad, CA) on 0.1µg/cm^2^ fibronectin coated tissue culture plates. Cells were single cell sorted for GFP into 96-well plates using a FACSAria II (Becton Dickinson, Franklin Lakes, NJ) cell sorter. Following expansion, viable cell clones were genotyped using digital droplet PCR (BioRad, Hercules, CA) with Taq-man specific fluorescent probes for the knock-in p.R183Q (referred as EC-R183Q), pseudo-WT (referred as EC-WT), and endogenous GNAQ alleles (IDT). The ddPCR primers and probes used in this study were designed using Primer3 Software (https://primer3.ut.ee/). Droplet generation and thermal cycling was conducted using standard BioRad ddPCR protocols. Briefly, the PCR mixture contained ddPCR Super Mix (BioRad), 0.25 mM of knock-in or reference probe, 0.9 mM of forward and reverse primer, and up to 30 ng of template DNA in 25µl total volume. The 20µl reaction mixture was emulsified into droplets using a QX100 or Automated Droplet Generator (BioRad) according to the manufacturer’s instructions. The droplets were then subject to the following thermal cycling profile: 10 minutes at 95°C, followed by 40 cycles of 30 seconds at 94°C, 60 seconds at 60°C annealing, 45 seconds at 72°C, and then kept at 4°C. Of note, during the 40 amplification cycles, the manufacturer suggested temperature ramp rate of 2°C/sec was used.

Samples were analyzed within 24 hours using the QX100 Droplet Reader (BioRad) and analyzed with QuantaSoft software (BioRad). Heterozygous mutant (i.e. a MUT: psuedoWT allele ratio of 1:1) and homozygous psuedoWT (i.e., only psuedoWT alleles detectable) were selected and expanded. ddPCR genotyping validation was done using standard Sanger Sequencing.

### Cell culture

The CRISPR edited cells designated EC-R183Q and EC-WT were expanded on 0.1µg/cm^2^ fibronectin (Millipore Sigma-Aldrich, FC010-10MG) coated plates at 15,000 to 20,000 cells/cm^2^ in endothelial cell growth medium-2 (EGM2, Lonza, CC3162), which contains endothelial cell basal medium-2 (EBM2), SingleQuot supplements (all except hydrocortisone and GA-1000), 10% heat-inactivated fetal bovine serum (FBS, Cytiva), and 1X glutamine-penicillin-streptomycin (GPS, Thermo Fisher). This growth media will be referred to as EGM2.

For experiments, the cells were grown in EBM2 without growth factors and with either 2% FBS or 0% FBS for 16-24 hours to promote quiescence, and throughout the remainder of the experiment as indicated.

### Western Blots

Cells were starved in EBM2 media with either 2% FBS or 0% FBS, no SingleQuots in either, for 24 hours. Cells were then lysed in a Pierce RIPA buffer (Fisher Scientific, PI89900) supplemented with PhosSTOP (Sigma Aldrich, 4906837001) and protease inhibitor cocktail (Sigma Aldrich, 11836170001). Cell lysate protein was quantified with the Bradford assay using Bio-Rad Protein Assay Dye Reagent (BioRad, 500-0006) and Pierce Bovine Serum Albumin Standard ampules (Thermo Fisher, 23209). Each SDS-PAGE gel lane was loaded with 15µg protein. Gels were transferred to polyvinylidene difluoride (PDVF) membranes, blocked in 5% bovine serum albumin/TBS-T for an hour or EveryBlot blocking buffer for 5 minutes and probed with primary antibodies (See Major Resources Table) overnight at 4°C with shaking. Antibodies against phosphorylated antigens were incubated in 5% bovine serum albumin/TBS-T, while antibodies against non-phosphorylated proteins were in 5% nonfat dry milk/TBS-T. Membranes were washed and probed with secondary isotype specific antibodies (See Major Resources Table) for 1 hour at room temperature. All signals were detected by enhanced chemiluminescence. The densiometric analysis was conducted using Fiji ImageJ or Image Studio software. Images of entire PDVF membrane for each Western Blot are shown in Supplemental Figures.

### Trans Endothelial Electrical Resistance (TEER) Assay

All TEER experiments were conducted following the same protocol unless noted otherwise. 30,000 cells/well were seeded on a fibronectin-coated (0.1mg/cm^2^) 96 well array with 10 interdigitated electrodes per well (Applied BioPhysics, 96W10idfPET ECIS array) in EGM2. Once cells adhered and formed a monolayer overnight (16hours), media was switched to reduced serum (2% FBS) EBM2 supplemented with thrombin (0.5 U/ml) (Sigma-Aldrich, 605190) to disrupt the nascent EC-EC contacts and follow subsequent barrier formation measured by electrical resistance (Ω). TEER was recorded after thrombin addition over 24 hours, with and without inhibitors. TEER was measured at multiple frequencies (62.5-64,000 Hz) in real-time using an ECIS^TM^ Zθ system (Applied BioPhysics). All ECIS measurements are reported in ohms (Ω) at 4,000 Hz, which is the optimal frequency for ECs. The percent change in resistance at 5 hours and 20 hours was quantified and normalized to appropriate control cells. 4 technical replicates for each condition were measured. Each experiment was repeated at least 3 independent times. A representative of the three is shown for each TEER experiment.

### siRNA experiments

EC-WT and EC-R183Q were transfected with siRNA using HiPerFect siRNA transfection reagent (Qiagen, 301705) following the manufacturer’s instruction.100nM final concentration GeneSolution siRNA targeted to ANGPT2, MAP2K1 and control non-target siRNA were used. Cells were analyzed by qPCR or by TEER 48 and 72 hours post transfection, respectively.

### shRNA experiments

EC-WT and EC-R183Q were transduced with ANGPT2 shRNA lentivirus or control lentivirus (Sigma) followed by 1µg/mL puromycin selection for 72 hours. Afterwards shANGPT2 and shControl cells were maintained in complete media without puromycin. Knockdown efficiency was confirmed by qPCR.

### qPCR

Total RNA was isolated using Qiagen RNeasy Mini Kit (Qiagen, 74104) according to the manufacturer’s protocol and was eluted in 15µl nuclease-free water. Isolated RNA concentration was verified using a NanoDrop 2000c spectrophotometer (Thermo Fisher Scientific). 1µg of total RNA was used for the reverse transcription using the iScript^TM^ cDNA Synthesis Kit (BioRad, 1708891). The cDNA was synthesized according to the manufacturer’s protocol (temperature cycles: 5 mins for 25°C (priming), 20 mins for 46°C (reverse transcription) and 1 min for 95°C (RT inactivation). RT-qPCR was performed using KAPA SYBR FAST Universal (KAPA Biosystems, KK4602). Amplification signals were detected on a QuantStudio 6 Flex System (Applied Biosystems). In brief, the PCR tubes (Applied Biosystems, Waltham, MA) were incubated at 95°C for 20sec before initiating the cycle for Taq polymerase activation. The cycling parameters were as follows: 95°C for 1sec; 60°C for 20sec with a melt curve as 95°C for 15sec; 60°C for 60sec. 2 technical replicates for each amplification reaction were performed.

The changes in cycle threshold (ΔCt) values were averaged and normalized with an endogenous housekeeping gene. 2^-ΔΔCt^ method was used to express relative gene expression.

Primer sequences are shown in the Major Resource Table. Primer amplification efficiencies were measured and pairs showing 90-95% efficiency were used.

### ELISA

Endothelial monolayers were washed with PBS twice, incubated in 2% serum media for 24 hours and supernatants were collected. ANGPT2 in the supernatants was measured with a commercial human Angiopoietin-2 Quantikine ELISA Kit (R&D systems, DANG20). An Angiopoietin-2 standard curve was run in parallel. Naïve serum was run as an additional control to normalize all the quantitative microplate reader values.

### Statistics

Data were analyzed and plotted by using GraphPad Prism v.10 (GraphPad Software). All the quantitative results are expressed as mean ± standard error mean (SEM). The differences between EC-WT and EC-R183Q were analyzed by 2-tailed Student t test. For experiments in which cells were treated with one inhibitor at multiple concentrations, the differences were assessed by Brown-Forsythe and Welch ANOVA test followed by Dunnett T3 multi comparison. For experiments in which cells were treated with siRNA or shRNA, two-way ANOVA was performed followed by Tukey’s multiple comparison test. Additional details about the statistics are stated in the figure legends. Significant differences were set at p<0.05.

## RESULTS

### Weakened endothelial barrier formation in CM and in p.R183Q endothelial cells

In 1977, it was hypothesized that cerebral vessels in SWS have abnormal vessel permeability^28^. Here and in our previous study we showed by Martius Scarlet Blue staining that CMs in the brain have increased extravascular fibrin suggesting permeability or leakiness^4^. Here we extend these findings to skin and brain CMs; both have increased extravascular fibrin compared to their non-CM controls (**Figure 1A-D**). Brain MRI studies with intravenous gadolinium contrast administration can provide further support for this finding giving indirect evidence of vascular permeability abnormalities in patients with SWS^29^. Longitudinal MRI on a SWS patient at 1 month and 15 months of age allows visualization of abnormally prominent and tortuous small vessels, coupled with abnormal diffuse contrast enhancement of the adjacent leptomeninges overlying the cortex seen at 15 months of age, which strongly suggests altered capillary permeability where the enhancement indirectly reflects leakage of gadolinium across the compromised blood-brain barrier, highlighting the developing functional abnormality of these vessels (**Figure 1E-H**).

**Figure 1.**
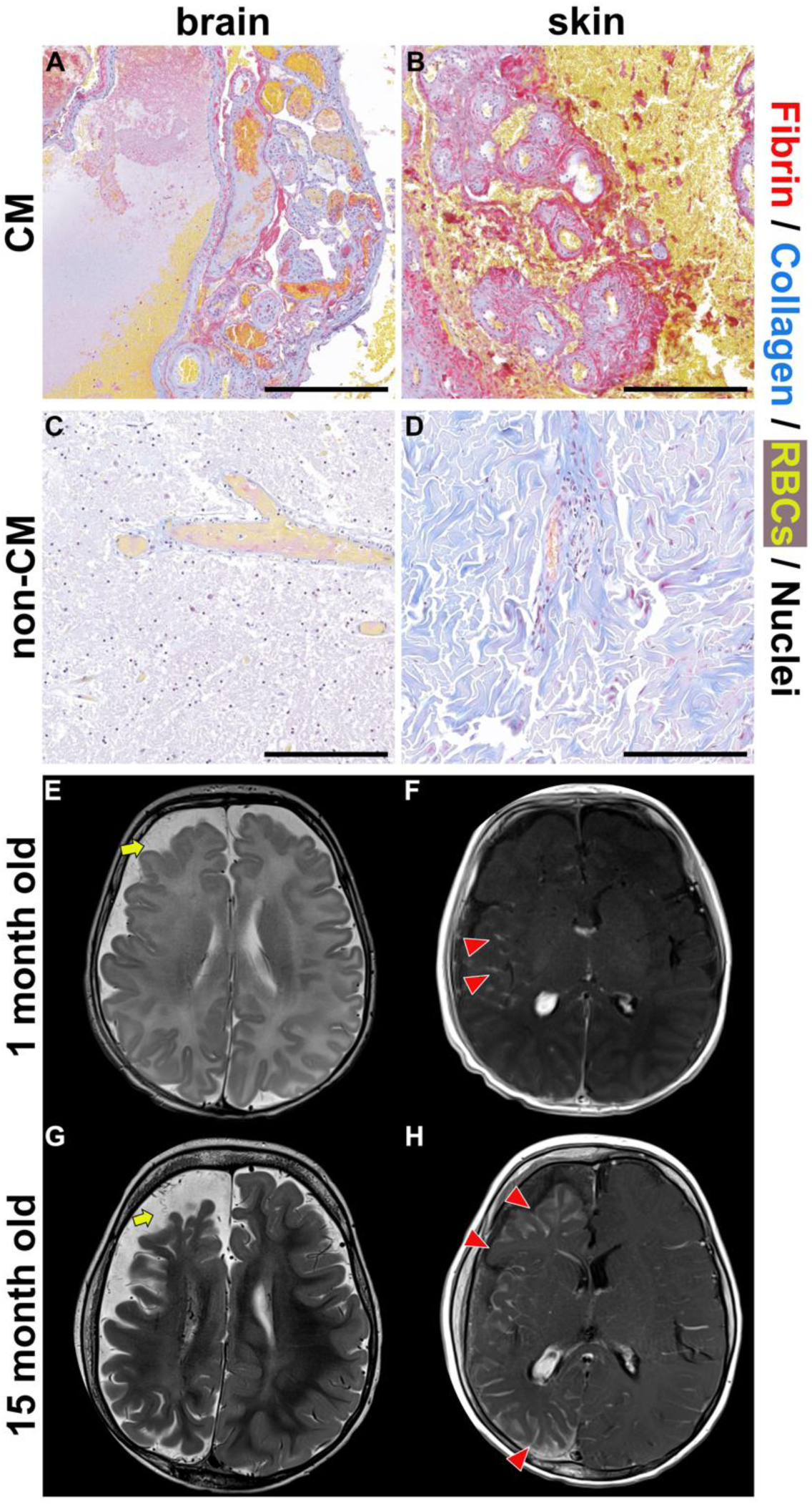
*Vascular permeability in Capillary Malformations*. Fibrin leakage in CM located in brain and skin. Martius Scarlet Blue staining of (**A**) CM brain (**B**) CM skin (**C**) non-CM brain (**D**) non-CM skin. Fibrin (red), collagen (blue), erythrocytes (yellow) and nuclei (blue/black). N=4 Scale bar=200μm. Brain MRI studies of SWS patient at 1 month and 15 months of age revealing progressive atrophy of the right cerebral hemisphere with associated progressive enlargement and tortuosity of subarachnoid small vessels (yellow arrows **E** and **G**) and associated progression of abnormal leptomeningeal enhancement along the cortical surface (red arrowheads **F** and **H**) suggestive of abnormal vascular permeability and regional disruption of the brain blood barrier.

To assess the vascular permeability and its causes, we used CRISPR-edited human ECs: EC- R183Q were edited to contain a single copy of the p.R183Q allele at the endogenous *GNAQ* locus and EC-WT were mock edited (**Figure 2A, Supplemental Figure 1**). We confirmed increased ANGPT2^19^ in the EC-R183Q versus EC-WT, grown in either reduced serum (2% FBS) (p=0.0448) or serum-free (0% FBS) (p=0.058). Gαq is shown for comparison; tubulin served as a loading control (**Figure 2B, Supplemental Figure 2C**). We measured permeability in the EC-R183Q and EC-WT using the TEER assay to quantify the EC-EC barrier formation. EC-R183Q displayed a sustained reduction (-36.54 ± 0.75%; p<0.0001) in TEER compared to EC-WT (**Figure 2D, E**). The results were similar when the TEER was conducted in 2% FBS (- 34.64 ± 0.55%) versus 0% (-44.43 ± 0.54%) FBS (**Supplemental Figure 3B, C**). Using a lentiviral p.R183Q ECs model^19^, we confirmed that the mutant ECs formed reduced barrier (- 15.47 ± 0.63%, p<0.0001) compared to the lentiviral EC-WT (**Supplemental Figure 4**). To verify the reduced barrier was a consequence of Gαq activity, we conducted the TEER experiment ± the Gαq inhibitor, YM254890 (100nM)^30-32^ (**Figure 2C**). YM254890 restored resistance (Ω) in EC-R183Q to EC-WT levels over the 26-hour period of TEER measurements (**Figure 2D**). YM254890 had no effect on the TEER in EC-WT (2.69± 0.44%), possibly because the EC-EC contacts were fully established (**Figure 2E**). YM254890 fully restored TEER in EC-R183Q (p=0.1458) to level seen in EC-WT at 20 hours (**Figure 2E**). This experiment establishes a role for the R183Q mutant Gαq in compromised endothelial barrier formation.

**Figure 2.**
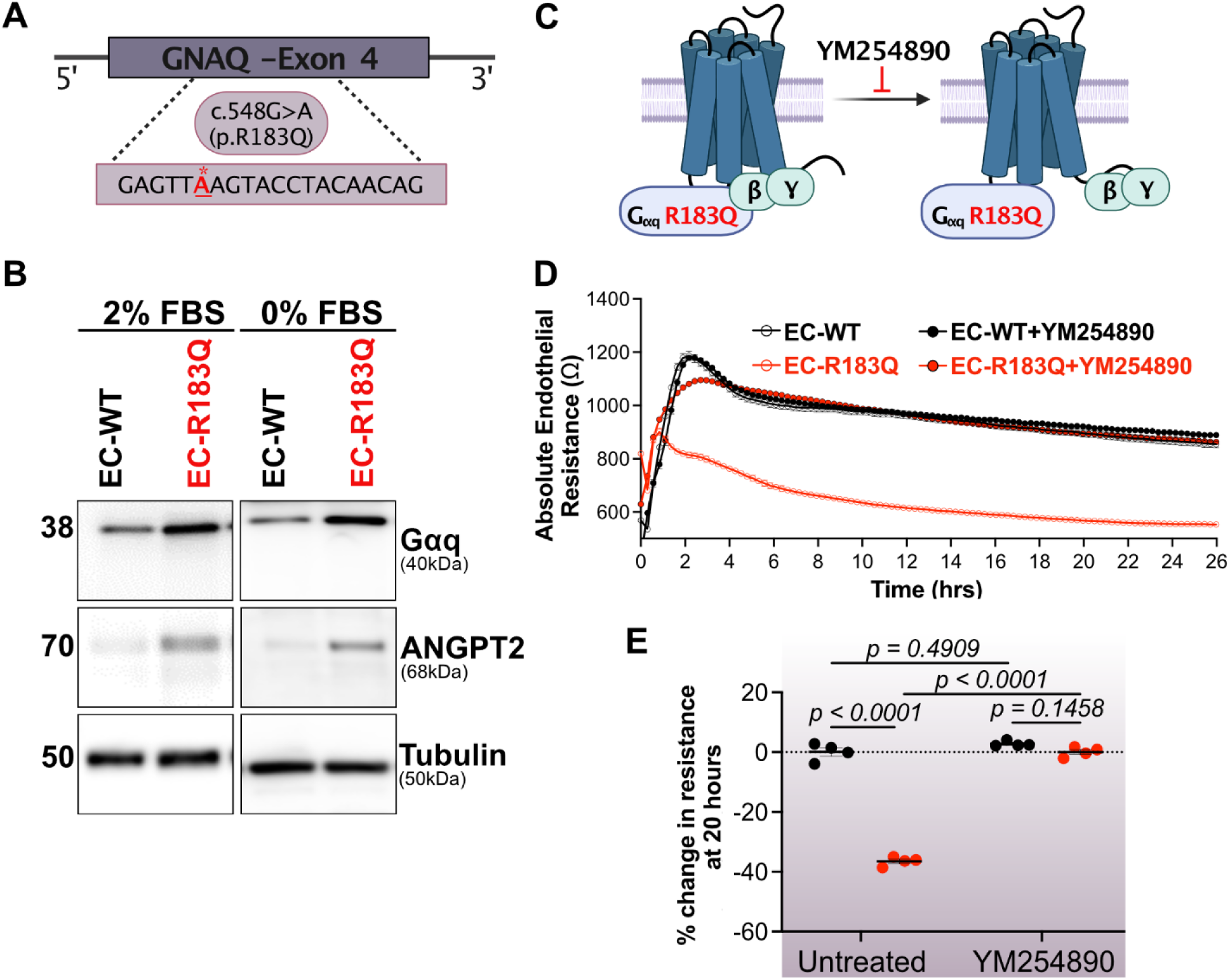
*GNAQ p.R183Q ECs form reduced barrier compared to wild-type ECs*. (**A**) Schematic of CRISPR editing to generate the p.R183Q mutation in ECs. (**B**) Western blot for Gαq (40kDa), angiopoietin2 (ANGPT2, 69kDa),and tubulin (50kDa) in EC-WT versus EC- R183Q in 2% FBS- (left) or 0% FBS-containing media (right) media for 24 hours. Full blots and their quantification are in Supplemental Figure 2. **C**) Schematic of Gαq dissociation from βγ subunits, blocked by the Gαq YM254890 inhibitor^32^. (**D**) Absolute endothelial resistance (Ω) in EC-WT (black open circles), EC-R183Q (red open circles), EC-WT treated with YM254890 (black closed circles), EC-R183Q treated with YM254890 (red closed circles) measured over 26 hours. (**E**) Quantification of percent change in resistance at 20 hours ± YM354890 (100nM). p- values were calculated by Brown-Forsythe and Welch ANOVA tests followed by Dunnett T3 multi comparison test. (**D,E**) n= 4 wells/condition were measured. 3 independent experiments were performed. A representative experiment shown.

### Modeling the somatic mosaicism in CM

ECs isolated from CM specimens present mutant allelic frequencies up to 21%, reflecting the mosaic nature of the *GNAQ* p.R183Q mutation in CM^15^. To assess the impact of mosaicism on endothelial permeability, we titrated EC-R183Q into f EC-WT cultures, ranging from 0-100%.

Increasing percentages of EC-R183Q reduced TEER over 24 hours (**Figure 3A**). Quantification at 20 hours showed that 10% EC-R183Q reduced TEER significantly (-14.78 ± 0.37; p<0.0001), suggesting that a small percentage of mutant ECs can have a detrimental effect on barrier formation (**Figure 3B**). Indeed, a trend towards increased permeability was seen at 5% EC- R183Q. We confirmed the mutant allelic frequency in the titration experiment by droplet digital PCR (**Supplemental Figure 5**). We next analyzed pERK (p=0.060) and ANGPT2 (p=0.0016) levels in cell lysates and found both increased in cultures with 50% EC-R183Q (**Figure 3C, D, E and Supplemental Figure 6**). We also determined ANGPT2 gene expression levels (**Figure 3F**) and secreted ANGPT2 measured by ELISA (**Figure 3G**). Both were significantly increased in cultures with 25% EC-R183Q.These results indicate that reduction in the mosaic endothelial barrier can occur before measurable increases in ANGPT2 or pERK (**Figure 3H**).

**Figure 3.**
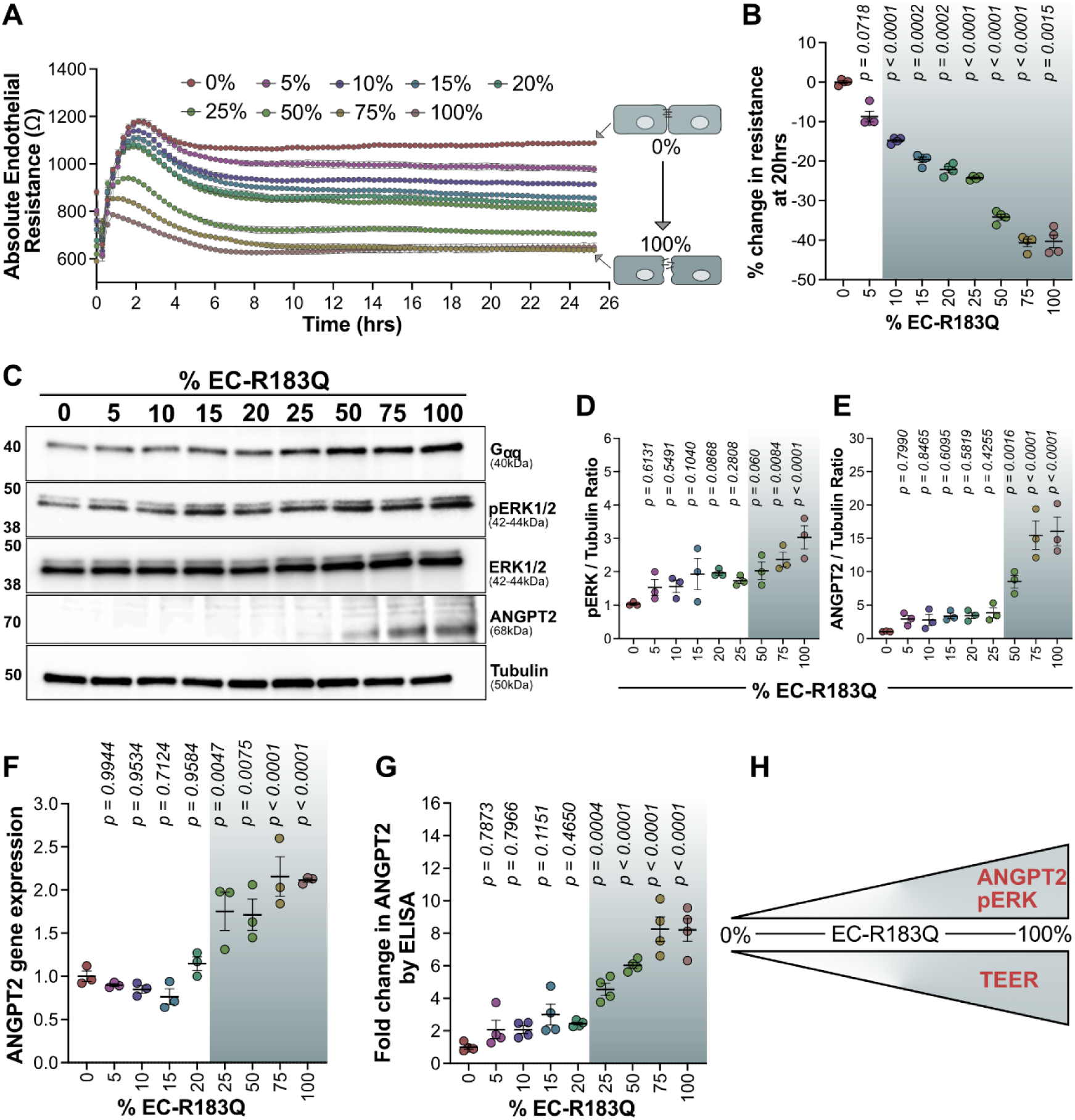
*Modeling p.R183Q mosaicism*. (**A**) Titration experiments were performed by adding EC-R183Q from 0-100% into EC-WT cultures. Absolute endothelial resistance (Ω) was measured over 26 hours. (**B**) Quantification of percent change in resistance at 20 hours. The p-values were calculated by Brown-Forsythe and Welch ANOVA test followed by Dunnett T3 multi comparison test. n= 4 wells/condition were measured. 3 independent experiments were performed. (**C**) WB for Gαq (40kDa), pERK1/2 (42- 22kDa), total ERK1/2 (42-22kDa) and ANGPT2 (68kDa) in EC-R183Q/EC-WT cultures. Tubulin (50kDa) served as loading control. (**D,E**) Western blot quantification of pERK1/2 and ANGPT2, normalized to tubulin, in C. The p-values were calculated by One-Way ANOVA test followed by Dunnett’s multiple comparison test. n=3. (**F**) ANGPT2 gene expression levels by qPCR at the end of TEER assay. The p-values were calculated by One-Way ANOVA test followed by Tukey’s multiple comparison test. n=4. 3 independent experiments were performed. (**G**) Secreted ANGPT2 levels were measured by ELISA at the end of TEER experiment. The p- values were calculated by One-Way ANOVA test followed by Tukey’s multiple comparison test. n=4. 3 independent experiments were performed. (**H**) Schematic showing increasing EC-R183Q coincides with increases in ANGPT2 and pERK and reduced TEER.

### MAPK pathway is involved in Gαq-R183Q endothelial permeability

PKC is one of the major effectors downstream of Gαq, and it is known to be involved in endothelial tight junctions^33^. Therefore, we tested a pan-PKC inhibitor, AEB071^24^, in the TEER assay (**Figure 4A**) and found significant rescue of TEER (23.9 ± 0.8%, p<0.0001) in EC-R183Q at 20 hours compared to untreated EC-R183Q (**Figure 4B**). One of the downstream targets of PKC is MAPK signaling, and thus we used an inhibitor of MEK, trametinib, which is currently used to treat fast flow vascular malformations^34-36^ as well as metastatic melanoma^37,38^. We measured the effects of increasing concentrations of (0, 0.1, 0.5, and1µM) over 24 hours in the TEER assay (**Figure 4C**). Significant yet partial changes were detected from 45.48 ± 3.37% to 40.98 ± 2.12% at 20 hours in EC-R83Q (**Figure 4D**). We next examined the effect of trametinib on the elevated ANGPT2 levels in EC-R183Q. Trametinib reduced ANGPT2 levels in a dose dependent manner, with 0.1μM trametinib causing significantly reduced ANGPT2 protein (p=0.0038) compared to untreated EC-R183Q. Similarly, 0.5μM (p=0.0002) and 1μM (p=0.0016) significantly reduced ANGPT2 levels (**Supplemental Figure 7A, B**). All three concentrations of trametinib reduced pERK significantly (p<0.001), as expected (**Supplemental Figure 7A, C**). The elevated level of Gαq in EC-R183Q was not significantly changed by increasing doses of trametinib (**Supplemental Figure 7**). Overall, these results suggest that MAPK pathway in EC- R183Q is involved in endothelial barrier formation and may contribute to elevated ANGPT2 levels in EC-R183Q. To confirm effects seen with trametinib, we tested two additional MAPK inhibitors: U0126, which selectively inhibits the activation of MEK1/2^39^ and RDEA119, a pharmacologic MEK inhibitor, also known as Refametinib^40^. U0126 (2.5uM) rescued the deficit in endothelial resistance in the TEER assay by 32.4 ± 3.29%, p=0.021) in EC-R183Q at 20 hours (**Supplemental Figure 8A, B**). RDEA119 (2.5μM) also rescued the endothelial resistance in the TEER assay by 29.2 ± 5.49%, p=0.0434) at 20 hours (**Supplemental Figure 8C, D**).

**Figure 4.**
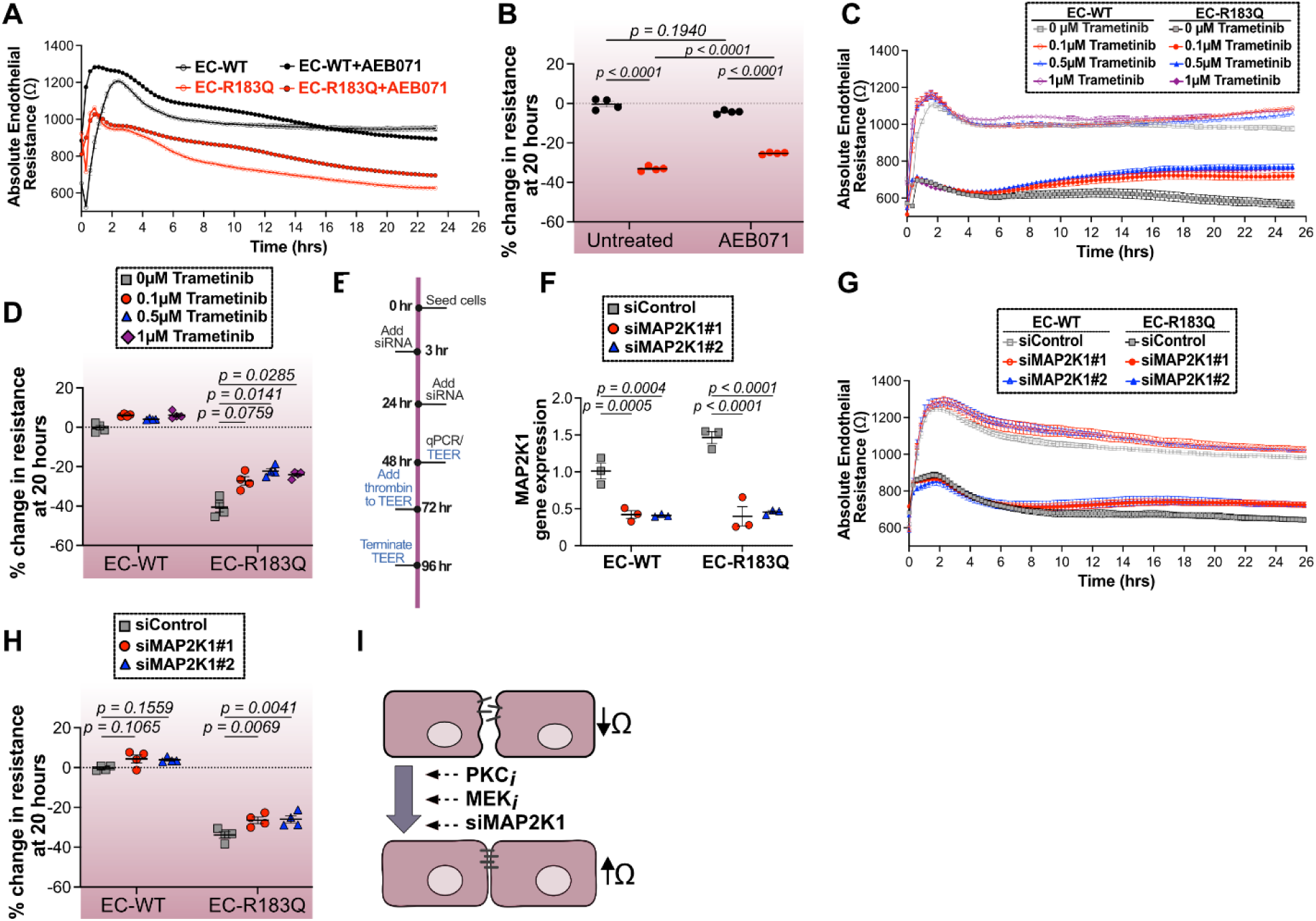
*PKC and MAPK pathways are involved in EC-R183Q barrier function*. (**A**) Absolute endothelial resistance (Ω) in EC-WT (black open circles), EC-R183Q (red open circles), EC-WT treated with AEB071 (black circles), and EC-R183Q treated with AEB071 (red circles) measured over 26 hours. (**B**) Quantification of percent change in resistance at 20 hours EC-WT and EC-R183Q ± AEB071 (10µM). The p-values were calculated by Brown-Forsythe and Welch ANOVA tests followed by Dunnett T3 multi comparison test. n= 4. 3 independent experiments were performed. (**C**) Absolute endothelial resistance (Ω) in EC-WT (open symbols) and EC-R183Q (closed symbols) ± increasing concentrations of trametinib (0 - 1µM) measured over 24 hours. (**D**) Quantification of percent change in resistance at 20 hours EC-WT and EC-R183Q ± Trametinib. The p-values were calculated by Brown-Forsythe and Welch ANOVA tests followed by Dunnett T3 multi comparison test. n= 4. 3 independent experiments were performed. (**E**) Schematic of MAP2K1 siRNA knock down experiment (**F**) MAP2K1 gene expression measured by qPCR 48 hours after siRNA transfection. (**G**) Absolute endothelial resistance (Ω) in EC-WT siControl (gray open squares), EC-WT siMAP2K1 (red open circles or blue open triangles), EC-R183Q siControl (gray closed squares), EC-R183Q siMAP2K1 (red closed circles or blue closed triangles) measured over 24 hours. (**H**) Quantification of percent change in resistance at 20 hours EC-WT and EC-R183Q ± siControl and two different siMAP2K1. The p-values were calculated by Brown-Forsythe and Welch ANOVA tests followed by Dunnett T3 multi comparison test. n= 4. 3 independent experiments were performed. (**I**) Summary of PKC and MAPK inhibitors that restored endothelial barrier function in EC-R183Q.

We next explored the consequences of MEK inhibition in EC-R183Q by knocking down MAP2K1 gene. We used different siMAP2K1 to knockdown MAP2K1 (**Figure 4E**) and achieved knockdowns of 72% and 69% (**Figure 4F**). Cells were treated with siMAP2K1 for 48 hours prior to measuring TEER (**Figure 4G**). siControl served as control in both cell types. Similar to trametinib, partial yet significant changes were observed for the two different siMAP2K1, which rescued the deficit in TEER by 21.9 ± 3.64%, p=0.0069 and 23.47 ± 5.76%, p=0.0041, respectively (**Figure 4H**). siMAP2K1 had no significant effect on TEER in EC-WT. These results suggest that both PKC and MAPK pathways are involved in reduced endothelial barrier in mutant R183Q ECs.

### ANGPT2 in Gαq-R183Q endothelial permeability

It has been previously reported that addition of ANGPT2 to primary murine brain ECs leads to decreased TEER compared to untreated ECs^41^. Since ANGPT2 is significantly increased in EC- R183Q, we tested its role in the reduced barrier formation observed in these cells. We first used siRNA to knockdown ANGPT2 (**Figure 5A**), with two successful knockdowns of 72% and 62% (**Figure 5B**). siANGPT2 knockdown or siControl was applied to EC-WT and EC-R183Q for 48 hours prior to measuring TEER (**Figure 5A**). The two siANGPT2 partially yet significantly rescued the deficit in TEER in the EC-R183Q by 50 ± 7.4%, p<0.0001 and 33 ± 1.66% p=0.0002, respectively) (**Figure 5C, D**). This finding was confirmed with shRNA lentiviral knockdown of ANGPT2 (**Figure 5E**); knockdown efficiency was determined by qPCR and found to be 54% **(Figure 5E, F**). Similar to siANGPT2, shANGPT2 partially yet significantly rescued the TEER deficit by 28.75 ± 2.86%, p<0.0001 in EC-R183Q at 20 hours (**Figure 5G, H**). Since the ELISA results indicated increased extracellular/secreted ANGPT2 in EC-R183Q (**Figure 3G**), we tested a function blocking antibody against human ANGPT2 (REGEN910)^42,43^ (**Figure 5I**). REGEN910 selectively binds to ANGPT2 and blocks its binding to the TIE2 receptor. We found REGEN910 had no effect on TEER in EC-WT or EC-R183Q (**Figure J, Supplemental Figure 9**). Quantification at 20 hours showed no significant difference in EC-R183Q treated with control IgG compared to EC-R183Q treated with REGEN910 (7.49 ± 3.03% p=0.3604) (**Figure K**). The activity of REGEN910 was verified in **Supplemental Figure 10** using phorbol 12- myristate 13-acetate treated normal human endothelial colony forming cells isolated from human cord blood^44^. Overall, these results show that knockdown of endogenous ANGPT2 resulted in increased TEER whereas addition of a function blocking anti-ANGPT2 had no effect. This suggests intracellular ANGPT2 may be required to diminish endothelial-endothelial barrier formation.

**Figure 5.**
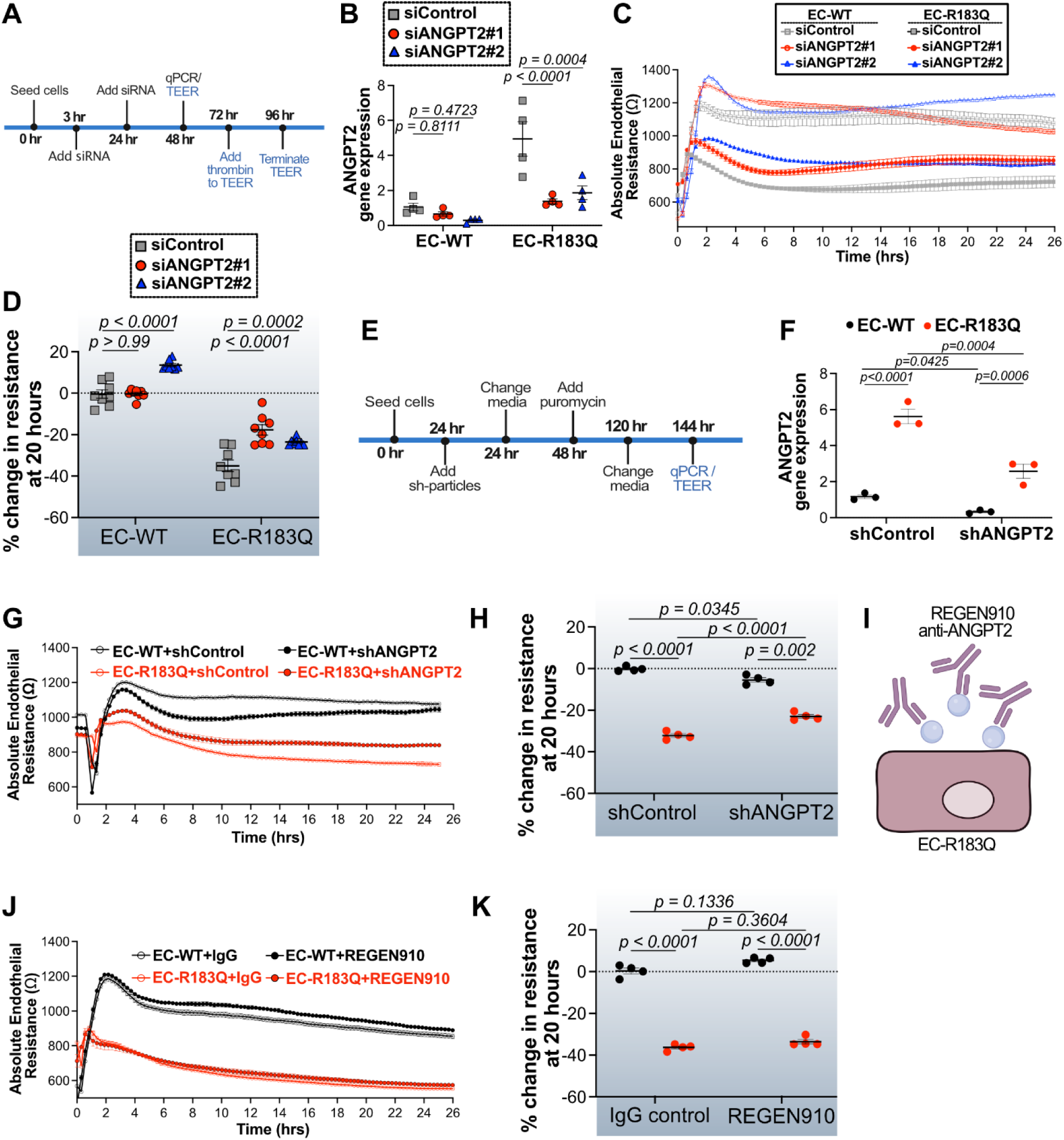
*ANGPT2 contributes to the reduced endothelial barrier in EC-R183Q*. (**A**) Schematic of siRNA knockdown experiment (**B**) ANGPT2 mRNA levels measured by qPCR at 48 hours after siRNA transfection. (**C**) Absolute endothelial resistance (Ω) in EC-WT siControl (gray open squares), EC-WT siANGPT2 (red open circles or blue open triangles), EC-R183Q siControl (gray closed squares), EC-R183Q siANGPT2 (red closed circles or blue closed triangles) measured over 26 hours. (**D**) Quantification of percent change in resistance at 20 hours EC-WT and EC-R183Q ± siControl and two different siANGPT2. The p-values were calculated by Brown-Forsythe and Welch ANOVA tests followed by Dunnett T3 multi comparison test. n= 8. 3 independent experiments were performed. (**E**) Schematic of shRNA knockdown experiment. (**F**) ANGPT2 gene expression levels by qPCR after shRNA transfection. (**G**) Absolute endothelial resistance (Ω) in EC-WT shControl (black open circles), EC-R183Q shControl (red open circles), EC-WT shANGPT2 (black closed circles), and EC-R183Q shANGPT2 (red closed circles) measured over 24 hours. (**H**) Quantification of percent change in resistance at 20 hours EC-WT and EC-R183Q ± siControl and shANGPT2. The p-values were calculated by Brown-Forsythe and Welch ANOVA test followed by Dunnett T3 multi comparison test. n= 4. 3 independent experiments were performed. (**I**) Schematic of anti-human ANGPT2 (REGEN910)^42,43^ addition to EC-R183Q. Schematic created in Biorender. (**J**) Absolute endothelial resistance (Ω) in EC-WT (black open circles), EC-R183Q (red open circles), EC-WT treated with REGEN910 (black closed circles), and EC-R183Q treated with REGEN910 (red closed circles) measured over 26 hours. (**K**) Quantification of percent change in resistance at 20 hours EC-WT and EC-R183Q ± REGEN910 (24nM). The p-values were calculated by Brown- Forsythe and Welch ANOVA test followed by Dunnett T3 multi comparison test. n= 4. 3 independent experiments were performed.

### Combination of ANGPT2 knockdown and trametinib effectively rescues reduced permeability phenotype in Gαq-R183Q endothelial cell

Since MEK inhibitors (trametinib, U0126, RDEA119) and silencing MEK partially increased TEER in EC-R183Q (**Figure 4, Supplemental Figure 8**) and knockdown of ANGPT2 had a similar effect (**Figure 5, Supplemental Figure 9, 10**), we hypothesized that combining the two would lead to a more complete rescue of endothelial barrier formation. To test this, EC-WT and EC-R183Q transduced with shANGPT2 were treated ± 1μM trametinib. shControl for both EC- WT and EC-R183Q were treated ± 1μM trametinib in parallel. Levels of cell associated ANGPT2 protein were evaluated by Western blot. We confirmed significant (p=0.0265) reduction of ANGPT2 protein levels with the knockdown (**Figure 6A, B, Supplemental Figure 11**). We next found shControl-EC-R183Q cells treated with trametinib to have noticeably reduced ANGPT2 (**Figure 6A, B, Supplemental Figure 11**). These results are consistent with **Supplemental Figure 7A** where we observed trametinib significantly reducing ANGPT2 levels in naïve EC- R183Q. These differences could potentially be from the manipulation of EC-R183Q with sh- lentiviral system. Importantly, shANGPT2 had no effect on pERK levels (**Figure 6A, C, Supplemental Figure 11**), which places MAPK upstream of ANGPT2. As expected, we confirmed that trametinib decreased pERK levels (**Figure 6C**). We next tested the shANGPT2- EC-R183Q ± trametinib in TEER assay (**Figure 6D**). EC-WT-shControl ± trametinib were used as control. shANGPT2-EC-R182Q treated with trametinib showed significantly restored TEER at 20 hours, with 67.45 ± 3.39%, p<0.0001 of the deficit erased, and TEER levels within 10.5 ± 1.55% of EC-WT (**Figure 6E**). In contrast, shANGPT2 alone or trametinib alone rescued about 1/3 of the deficit in EC-R183Q (**Figure 6E**). Overall, these results suggest that the combination of ANGPT2 knockdown and trametinib nearly restores the barrier function in an additive manner Using telomerase immortalized microvascular Gαq-EC-R183Q cells, Zecchin and colleagues showed that there is active calcium influx from outside the cells in the mutant ECs^45^. This was verified using the CRAC channel inhibitor, CM4620^45^. We tested CM4620 in the TEER assay and found no effect at concentrations 0.125-1μM on the endothelial barrier in EC-R183Q or EC- WT (**Supplemental Figure 12**) suggesting that calcium flux into the cytoplasm does not affect endothelial barrier formation. We also tested Tacrolimus (200nM), an inhibitor of calcineurin pathway and found no change in the TEER assay measurements (**Supplemental Figure 13**).

**Figure 6.**
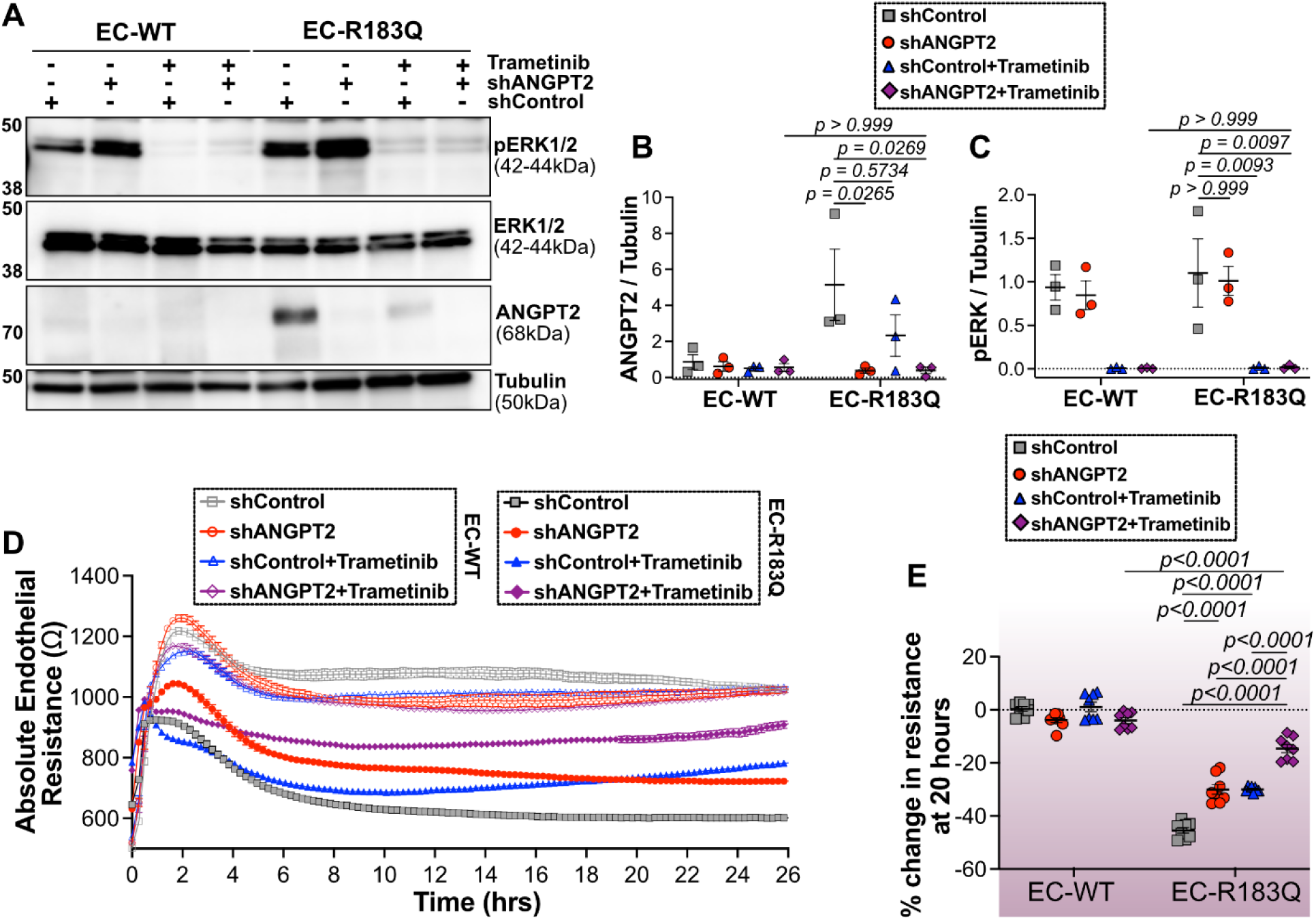
*Combined ANGPT2 and MAPK inhibition restores EC-R183Q endothelial barrier*. (**A**) pERK1/2 (42-22kDa), total ERK1/2 (42-22kDa) and ANGPT2 (68kDa) in EC-WT and EC- R183Q treated with shControl, shANGPT2, shControl treated with Trametinib, and shANGPT2 treated with Trametinib were detected by WB. Tubulin served as loading control. WB quantification of (**B**) ANGPT2 and (**C**) pERK1/2 normalized to tubulin. The p-values were calculated by Two-Way ANOVA test followed by Šídák’s multiple comparison test. n=3. (**D**) Absolute endothelial resistance (Ω) in EC-WT (open symbols) and EC-R183Q (closed symbols) ± treated with shControl (gray squares), shANGPT2 (red circles), shControl treated with Trametinib (blue triangles), shANGPT2 treated with Trametinib (purple diamonds) measured over 24 hours. (**E**) Quantification of percent change in resistance at 20 hours EC-WT and EC- R183Q ± treated with shControl (gray squares), shANGPT2 (red circles), shControl treated with Trametinib (blue triangles), shANGPT2 treated with Trametinib (purple diamonds). The p-values were calculated by Brown-Forsythe and Welch ANOVA tests followed by Dunnett T3 multi comparison test. n= 8. 3 independent experiments were performed.

These two findings suggest that calcium and the calcineurin pathway play minimal roles in the reduced endothelial-endothelial barrier in EC-R183Q. Moreover, we tested the mTOR inhibitors everolimus and rapamycin in our TEER assay and found no effects on the on EC-R183Q barrier function (**Supplemental Figure 14**).

## DISCUSSION

In this study, we show that *GNAQ* p.R183Q expressing ECs (EC-R183Q) form an endothelial monolayer with increased permeability, indicated by reduced TEER, compared to EC-WT. Furthermore, we show that a small percentage of mutant EC in the endothelial monolayer increased permeability in an outsized manner, which may be relevant to the mosaic nature of CM vessels. Genetic knockdown and pharmacologic inhibition experiments show that ANGPT2 and MEK1 contribute to reduced endothelial barrier formation in the EC-R183Q (**Figure 7**). We posit that the weakened endothelial barriers formed in the presence of EC-R183Q mirrors the extravascular fibrin seen in histological sections of CM and the apparent permeability seen by contrast MRI in a child with SWS.

**Figure 7.**
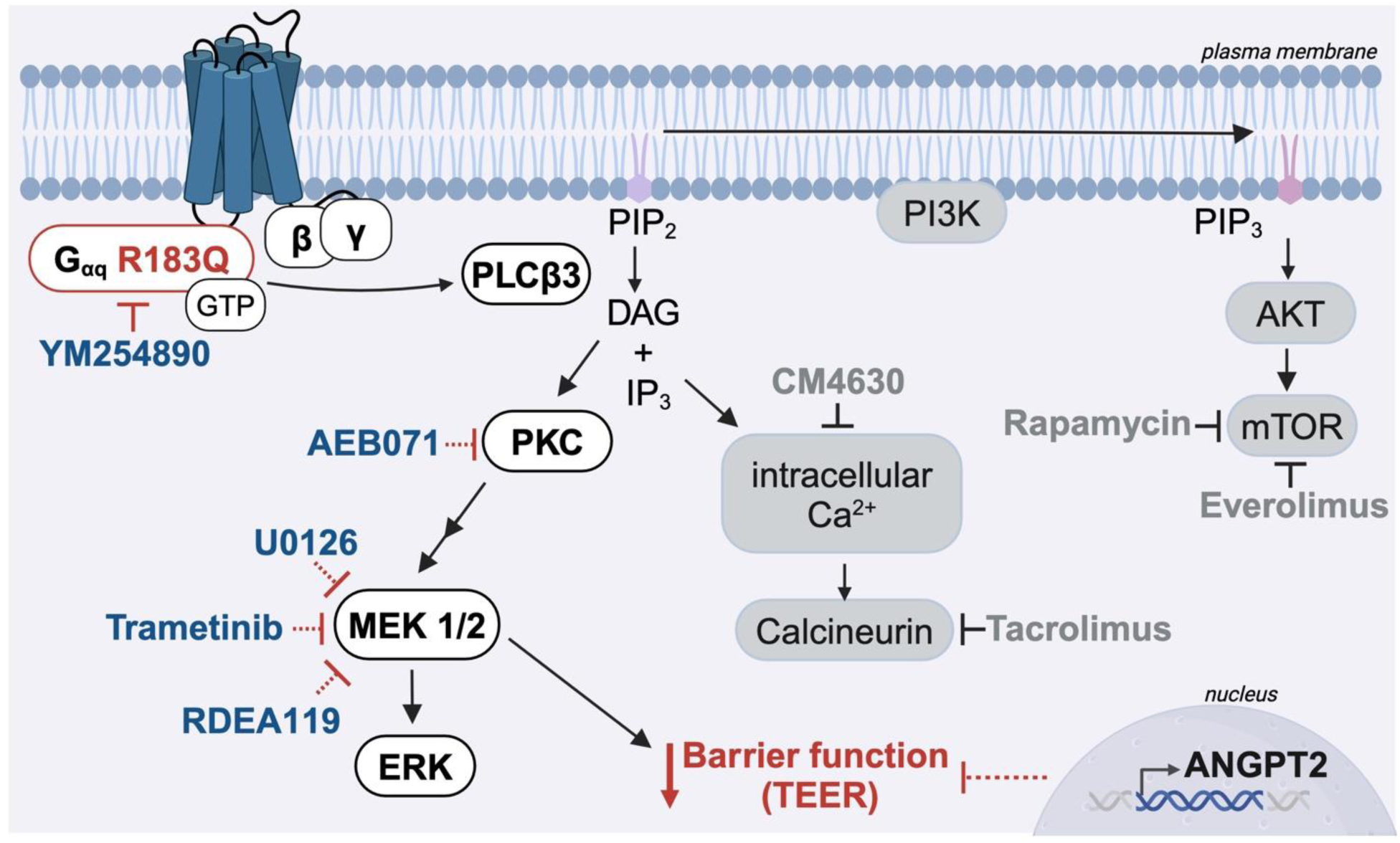
*Summary of the Gaq-R183Q pathway and role of downstream effectors in endothelial barrier function.* Inhibitors with no effect on Gαq-R183Q endothelial barrier function shown in gray. Schematic created in Biorender.

Permeability abnormalities of the endothelium can be qualitatively and quantitatively assessed through brain MRI studies with intravenous gadolinium contrast administration. The characterization of abnormal leptomeningeal contrast enhancement in the affected regions of the brain serves as a strong qualitative indicator of altered capillary permeability, reflecting a compromised blood-brain barrier^29^. In future studies, the degree of abnormal permeability could be further evaluated using advanced sequences where capillary permeability measurements (Ktrans) can be obtained using dynamic contrast-enhanced magnetic resonance perfusion^46-48^. This technique can measure the accumulation of gadolinium-based contrast agent in the extravascular-extracellular space, which can be seen in different conditions affecting the brain, including tumors, vascular malformations, infections and different inflammatory processes.

In addition to human studies, permeability in affected blood vessels has been described in one of the first animal models of GNAQ p.R183Q capillary malformation^26^. In this mouse model, perfusion of Evans blue dye showed severe leakage and irregular expression of claudin-5 in the leptomeningeal vessels in the mutant mice. This is consistent with the lack of junctional markers (zona occludin1 and claudin-5) we previously observed in CM brain and skin blood vessels^27^..

Vascular permeability has also been shown in other vascular malformations such as cerebral cavernous malformation (CCMs)^49,50^, and arteriovenous malformation^51-53^. Particularly in CCMs, knockdown of CCM3 in brain ECs was shown to reduce TEER *in vitro* and tight junction stability *in vivo*^54,55^.

Our experiments show that ANGPT2 and MAP2K1 contribute to the increased permeability in EC-R183Q monolayers. Angiopoietins are crucial angiogenic factors that play fundamental roles in vascular development and angiogenic processes. Angiopoietin-1 binds to the receptor tyrosine kinase TIE2 to promote vessel formation and stabilization whereas ANGPT2 in most settings antagonizes TIE2 signaling resulting in vessel destabilization, remodeling, and angiogenesis. Upregulation of ANGPT2 occurs in many types of advanced tumors^56-61^ and vascular anomalies^19,62-65^. ANGPT2 has also been proposed as a serum biomarker for lymphatic anomalies^66^ and has been associated with blood brain barrier leakiness in early Alzheimer’s disease^67^. In another type of vascular malformation, cerebral CCMs exhibit brain hyperpermeability, cerebral hemorrhage, seizures, and stroke^50,68^. CCM3 loss of function results in elevated ANGPT2 in ECs. A neutralizing antibody against ANGPT2 was shown to normalize defects caused by CCM3 deficiency^64^. In another study associated with cerebrovascular leakage and brain edema, ANGPT2 was shown to mediate blood brain barrier permeability via endothelial paracellular and transcellular routes^41^. Overall, ANGPT2 is reported to be elevated in many vascular malformations and antibodies against it have been pre-clinically used to stabilize vessel permeability. Its role in EC-R183Q barrier formation suggests targeting ANGPT2 in CM would promote vascular integrity and prevent leakage.

Trametinib is a MEK1/2 inhibitor approved for certain cancers and increasingly explored for vascular malformations linked to MAPK pathway hyperactivation. Trametinib selectively blocks phosphorylation and activation of ERK1/2, thereby reducing downstream signaling needed for cell proliferation, survival, and angiogenesis. Trametinib is currently in phase 2 clinical trial for arteriovenous malformation. In KRAS-mutant EC models, trametinib reduced arteriovenous malformation size and normalized vessel morphology^34^.

Sirolimus, a mammalian target of rapamycin (mTOR) inhibitor, has been tested in in SWS patients for improvements in cognitive functions and topically for improvement in pulse dye laser treatment of skin CMs. Sirolimus was well tolerated by adult SWS in a phase 2/3 clinical trial^69^. Sirolimus combined with pulsed dye laser for patients with port wine birthmark showed reduced skin pigmentation and lower number of vessels in histologic analysis^70^. In our *in vitro* TEER model, we tested two mTOR inhibitors, everolimus and sirolimus (aka rapamycin), for effects on EC-R183Q barrier formation in the TEER assay. Neither mTOR inhibitor influenced the endothelial barrier formation. Sirolimus may affect other properties of the *GNAQ*-R183Q endothelium, which could be explored in future studies.

There are several limitations in our study. For most of the *in vitro* experiments we used telomerase-expressing ECs. The cells express endothelial markers as expected, proliferate well in culture, and form blood vessels when xenografted into immune deficient mice (not shown), however the telomerase ECs may not fully represent the growth and functional properties of ECs from CMs in the skin and brain wherein somatic *GNAQ* R183Q mutation is present. A second limitation is that the knockdown of ANGPT2 in EC-R183Q was not 100% such that remaining ANGPT2 might have affected some results. Third, experiments to examine potential paracrine signaling from neighboring cells, for example by incorporating mural cells into the experimental design, would be informative.

In conclusion, our study demonstrates that Gαq-R183Q expressing ECs form reduced endothelial barrier in the TEER assay. Signs of increased vascular permeability are evident in human studies of CM and SWS, which strengthens the relevance of our findings using ECs in cell culture. Histopathology shows that there is extravascular fibrin in both brain and skin CMs and longitudinal MRI of the brain of a child with SWS indicates capillary permeability in the leptomeninges develops during infancy. Additional studies are needed as this is a n=1 case. A murine developmental model of brain CM also shows increased permeability in brain vessels of mice with inducible expression of GNAQ pR183Q^26^, which further demonstrates endothelial permeability as a pathologic feature of CM. Our discovery of increased permeability in human ECs expressing the mutant Gαq provides a platform for drug testing that we hope will address clinical needs of these patients.

## ACKNOWLEDGEMENTS

The authors would like to thank the Vascular Biology Program Microscopy Imaging Core, and the Histology Core at Boston Children’s Hospital. We would like to thank Drs. Raj Gupta and Gavin Schnitz for donating the teloHAECs used for CRISPR editing and subsequent experiments. We would also like to thank the Biopolymers Facility at Harvard Medical School for assistance with generating and characterizing the mutant teloHAECs cell lines.

## AUTHORS CONTRIBUTIONS

SN conceived the project, collected data and material, designed, and executed experiments, and wrote the manuscript. JB supervised the project and wrote the manuscript. MB assisted with analyzing the TEER data under SN’s supervision, JWS performed the western blots and its quantification. MV generated the GNAQ cell model. PS, LM, YSC assisted MV with the GNAQ cell model generation under AKG’s supervision. All authors read the manuscript and approved.

## SOURCES OF FUNDING

Research reported in this manuscript was supported by the National Heart, Lung, and Blood Institute, part of the National Institutes of Health, under Award Number 5R01HL127030 (J.B., A.K.G). S.N. was supported by F32HL172637 and is currently supported by K99 Career Development Award - K99HL177326. M.V. was supported by F32HD107878. We also extend gratitude and thanks to Taylor/McDonald Family. The content is solely the responsibility of the authors and does not necessarily represent the official views of the National Institutes of Health.

## DISCLOSURES

None

## Notes

### Competing Interest Statement

The authors have declared no competing interest.

