## Supplemental Figures 1-14 for "MAPK Signaling and Angiopoietin-2 Contribute to Endothelial Permeability in Capillary Malformations"

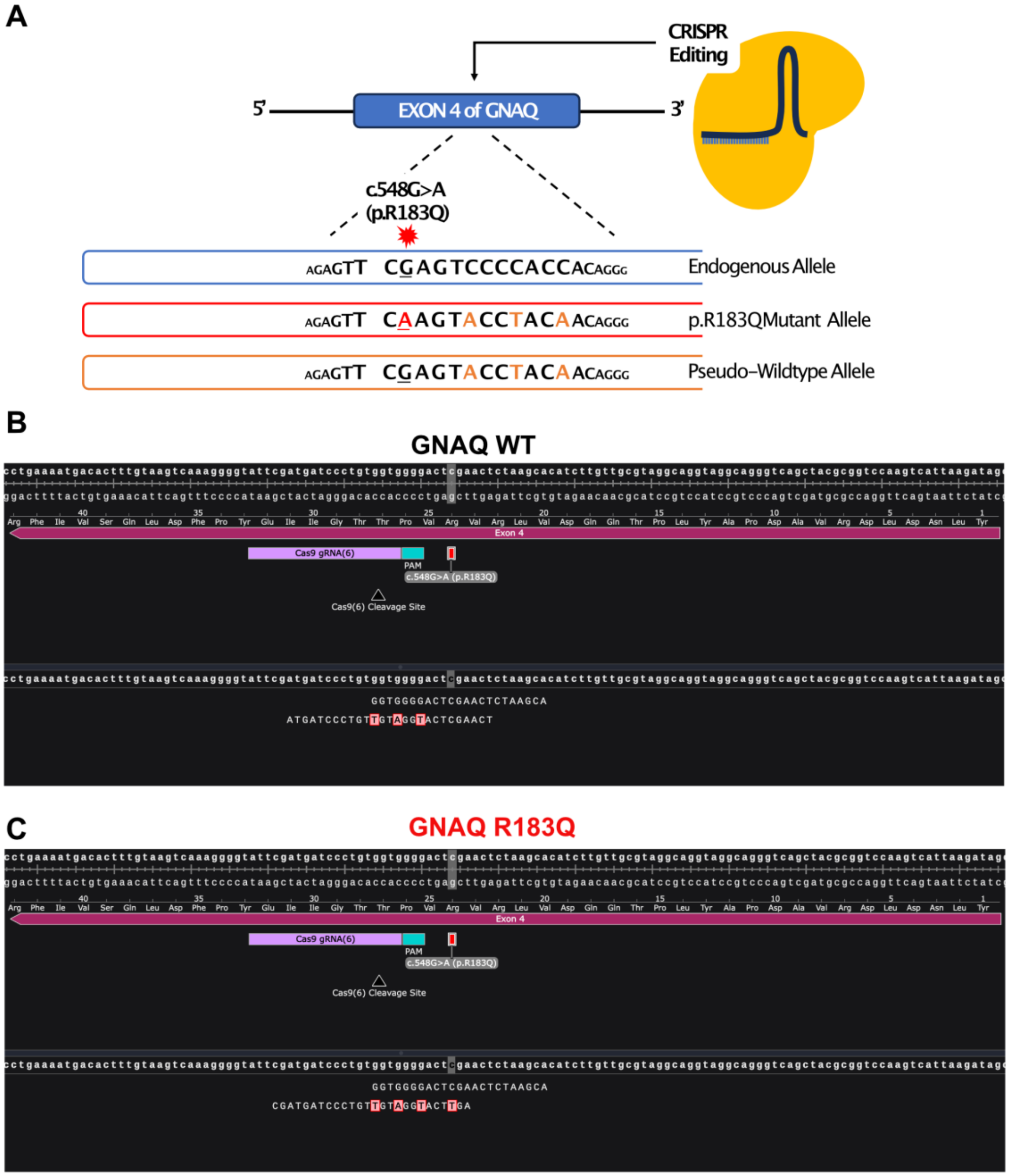


*Supplemental Figure 1 – Telomerase expressing human aortic endothelial cells (teloHAECs) with GNAQ R183Q mutation.* (**A**) Schematic of the location of mutation insertion. Orange bases are synonymous SNVs to avoid rebinding of Cas9 and additional cleavage. GNAQ sequences in (**B**) EC-WT and (**C**) EC-R183Q.

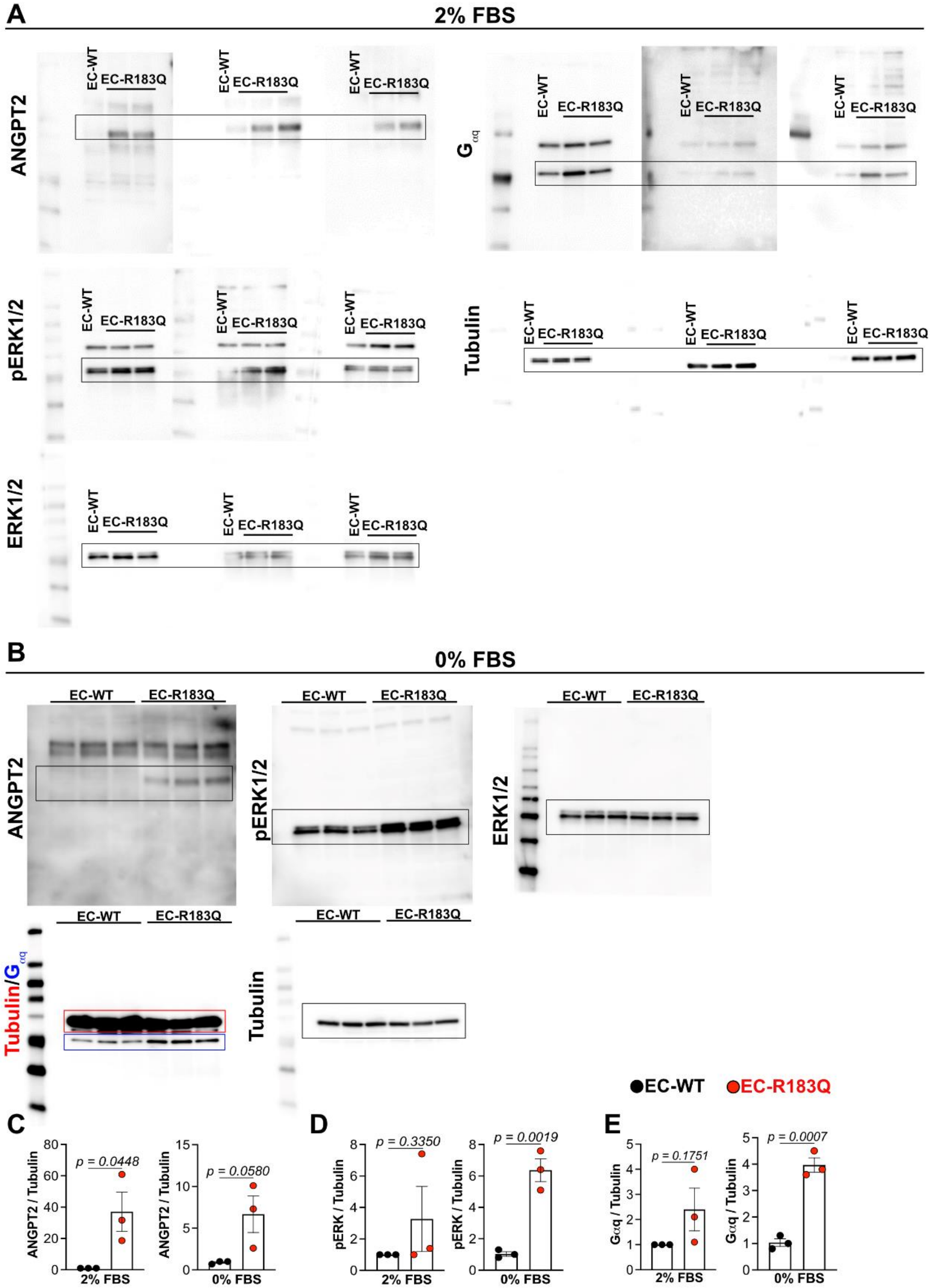


*Supplemental Figure 2 – Uncut WBs from EC-WT and EC-R183Q cultured in 2% or 0% FBS-containing media for 24 hours.* (**A**) 2% FBS (**B**) 0% FBS WBs probed with anti-ANGPT2, anti-pERK1/2, anti-ERK1/2, anti-Tubulin, or anti-Gαq. Quantification of WB bands detected in cells in 2% FBS and 0% FBS for (**C**) ANGPT2 (**D**) pERK and (**E**) Gαq.

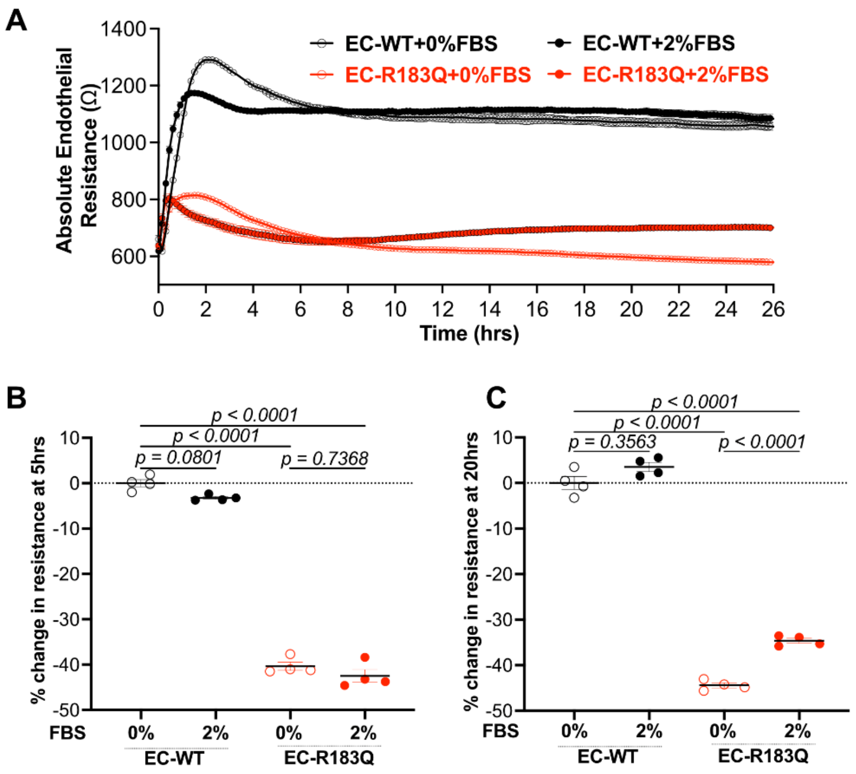


*Supplemental Figure 3 – TEER measurements on EC-WT and EC-R183Q cultured in media with 0% or 2% FBS, both without growth factors.* (**A**) Absolute endothelial resistance (Ω) of EC-WT (black) and EC-R183Q (red) in cultures with 0% or 2% FBS over 24 hours. Quantification of percent change in resistance at (**B**) 5 hours and (**C**) at 20 hours. n= 4. 3 independent experiments were performed. The p-values were calculated by brown-Forsythe and Welch ANOVA test followed by Dunnett T3 multiple comparison test.

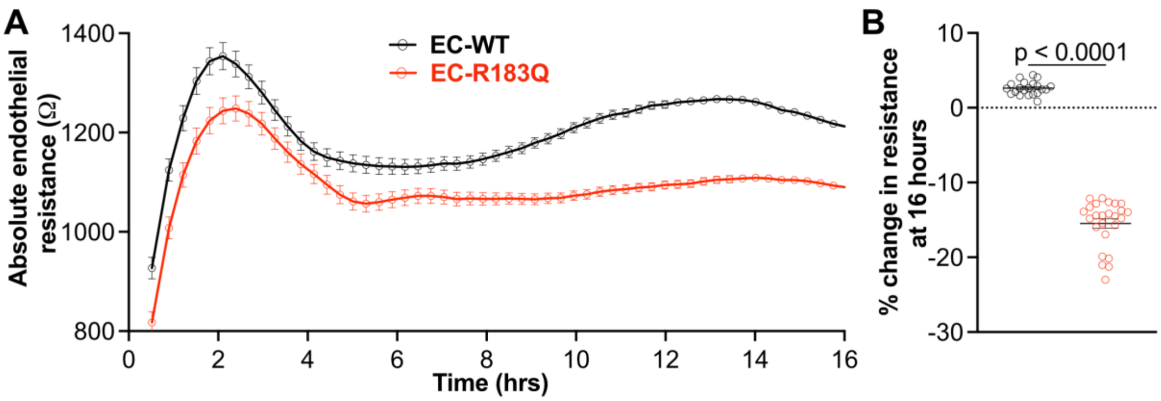


*Supplemental Figure 4 – Lentivirally expressed GNAQ R183Q in human endothelial colony forming cells results in reduced TEER compared to lentivirally expressed GNAQ WT in human endothelial colony forming cells*[4]*.* (**A**) Absolute endothelial resistance (Ω) of lentivirus transduced EC-WT (black) and EC- R183Q (red) over the period of 16 hours (**B**) Percent change in resistance at 16-hour time point. The p-value was calculated by two tailed student t-test. n=16.

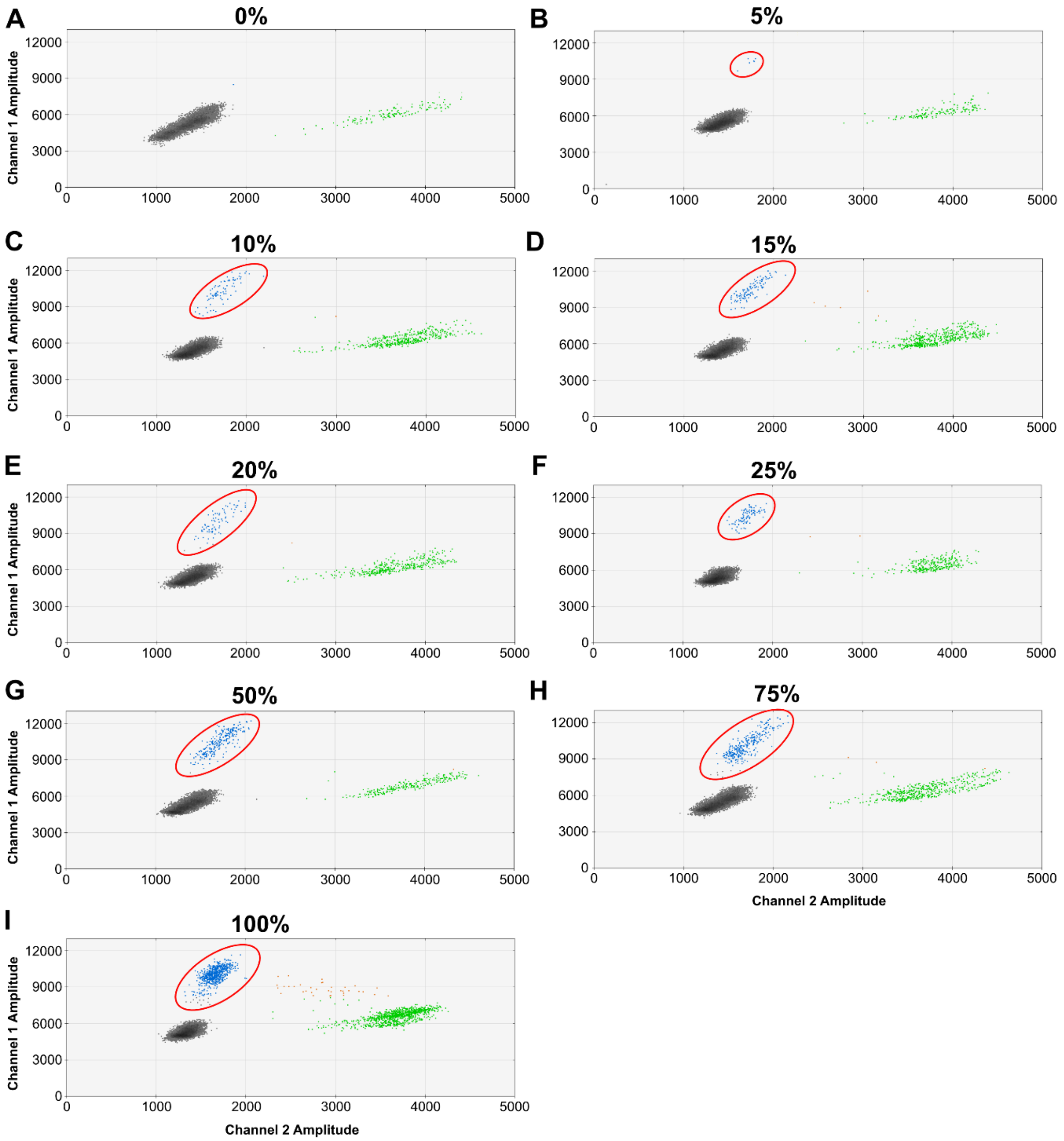


*Supplemental Figure 5 – Mutant allelic frequency measured by droplet digital PCR in cultures with increasing percentages of EC-R183Q.* ddPCR was performed to verify mutant allelic frequency in the titration experiment. (**A**) 0% (**B**) 5% (**C**) 10% (**D**) 15% (**E**) 20% (**F**) 25% (**G**) 50% (**H**) 75% (**I**) 100% EC-R183Q. Channel 1 – GNAQ Mutant R183Q and Channel 2 – GNAQ WT. N=3. 3 independent experiments were performed. Representative images are from one experiment. Red circles indicate the mutant allele.

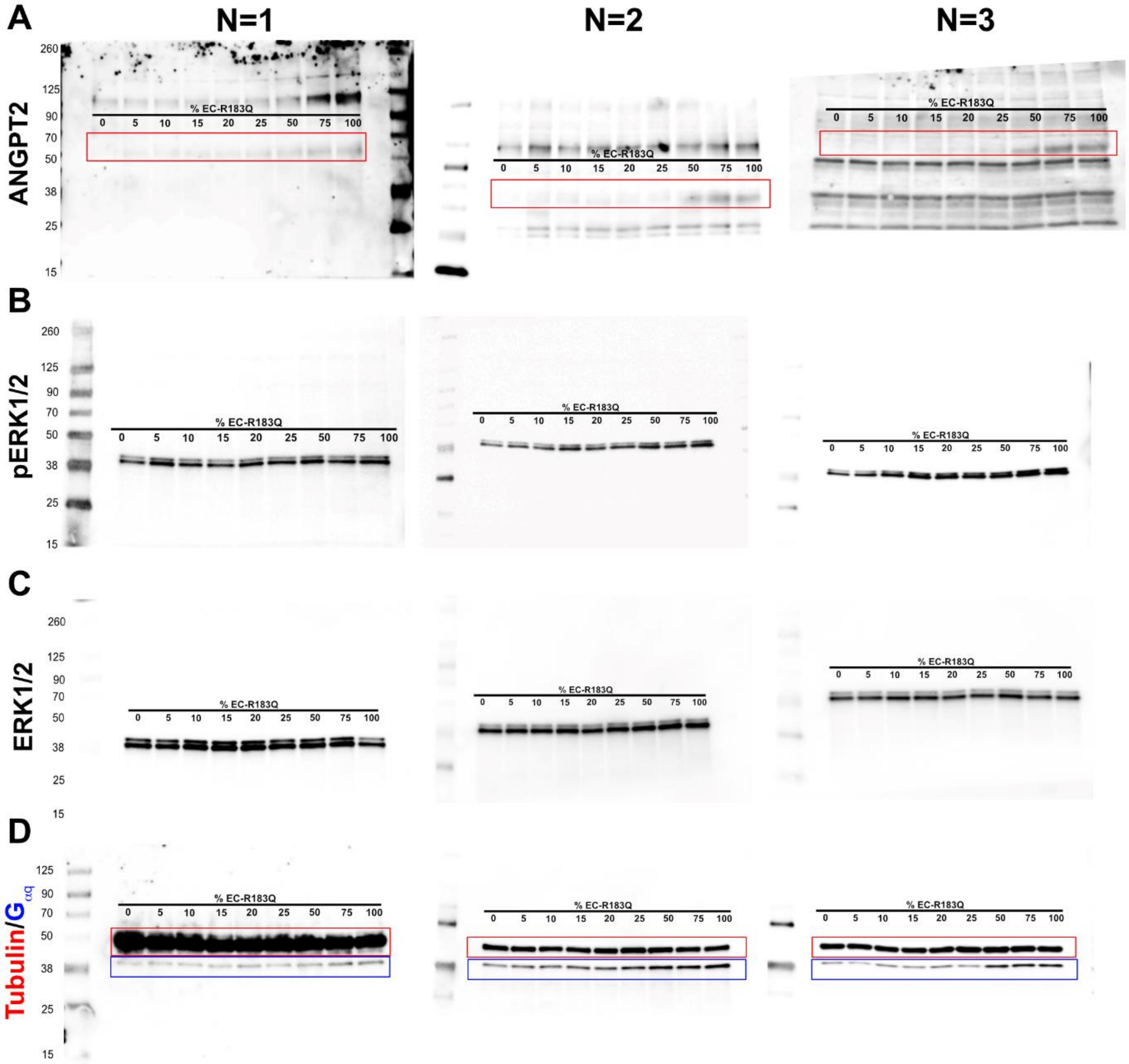


*Supplemental Figure 6 – Uncut WBs from 3 independent titration experiments.*
WBs probed with (**A**) anti-ANGPT2 (**B**) anti-pERK1/2 (**C**) anti-ERK1/2. (**D**) anti-tubulin and anti-$G_{\alpha q}$. The catalog numbers and details for each antibody are provided in the *Major Resource Table*. Molecular weight markers are shown on the left side of each membrane.

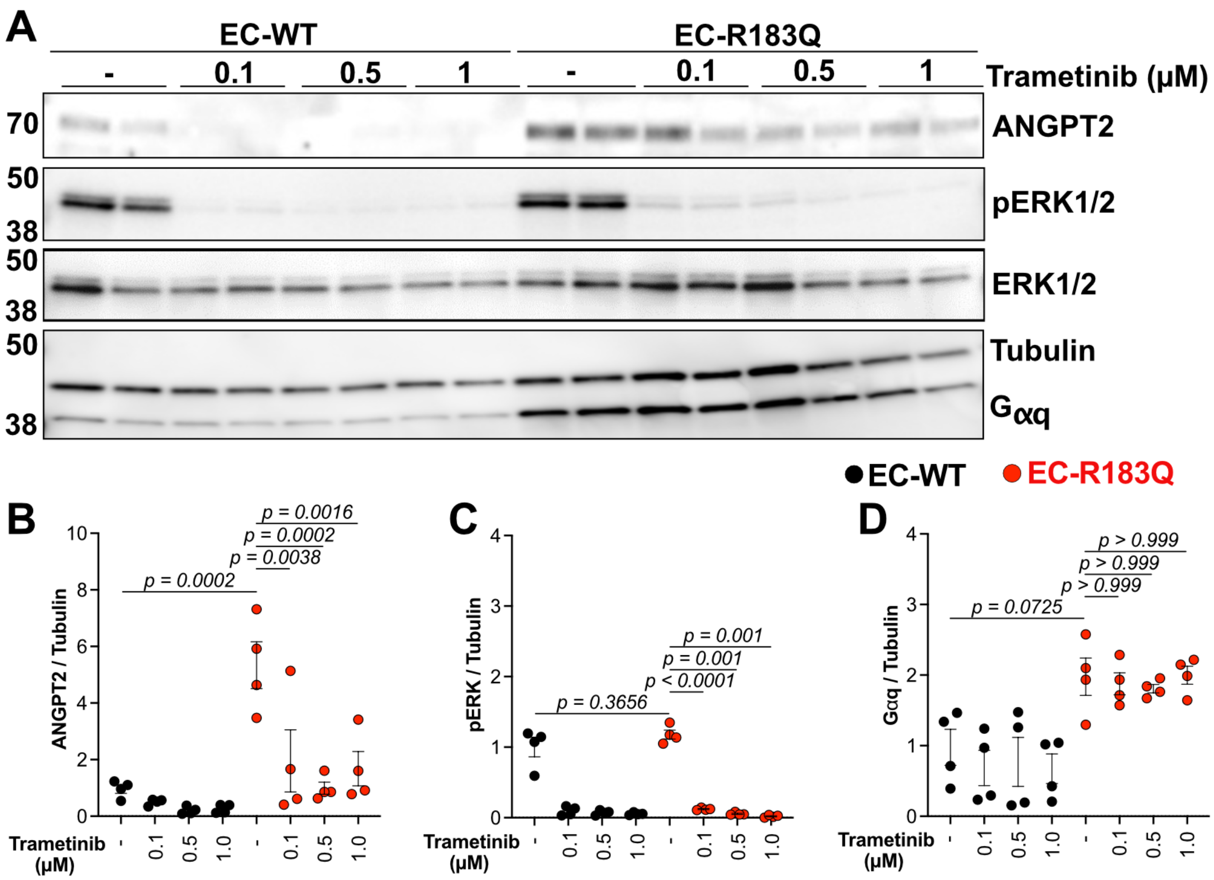


*Supplemental Figure 7 – Trametinib reduces ANGPT2 and pERK levels.*

(**A**) ANGPT2, pERK1/2, total ERK1/2 and Gαq were detected by WB of cell lysates from EC-WT and EC-R183Q ± three different concentrations of Trametinib. Tubulin served as loading control. WB quantification of (**B**) ANGPT2 and (**C**) pERK1/2 (**D**) $G\alpha_q$ normalized to tubulin. The p-values were calculated by One-Way ANOVA test followed by Tukey's multiple comparison test. n=3.

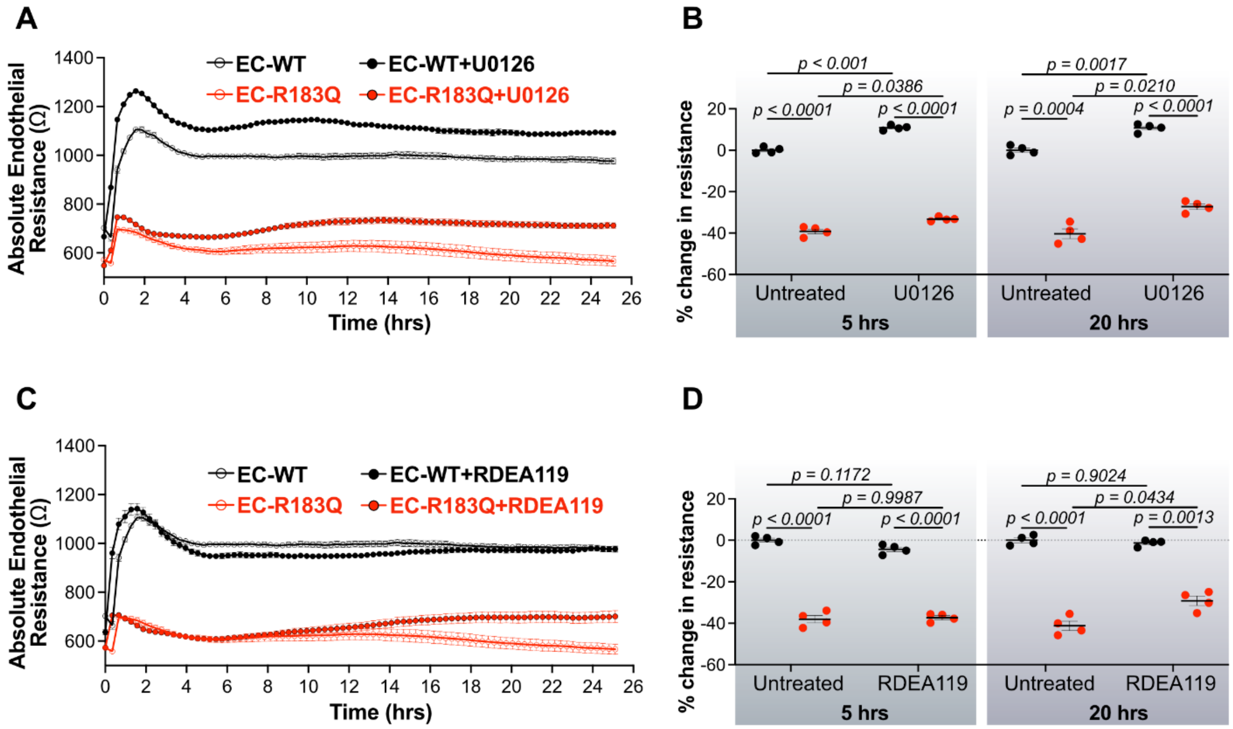


*Supplemental Figure 8 – MEK inhibitors partially restore endothelial barrier in mutant GNAQ R183Q.* (**A**) Absolute endothelial resistance (Ω) of EC-WT (black open circle), EC-R183Q (red open circles), EC-WT treated with U0126 (black closed circles), EC-R183Q treated with U0126 (red closed circles) over 24 hours. (**B**) Quantification of percentage change in resistance at 5 hours and 20 hours EC-WT and EC-R183Q ± U0126 (2.5µM). (**C**) Absolute endothelial resistance (Ω) of EC-WT (black open circle), EC-R183Q (red open circles), EC-WT treated with RDEA119 (black closed circles), EC-R183Q treated with RDEA119 (red closed circles) over 24 hours. (**D**) Quantification of percent change in resistance at 5 hours and 20 hours EC-WT and EC-R183Q ± RDEA119 (2.5µM). n= 4. 3 independent experiments were performed. The p-values were calculated by Brown-Forsythe and Welch ANOVA test followed by Dunnett T3 multiple comparison test.

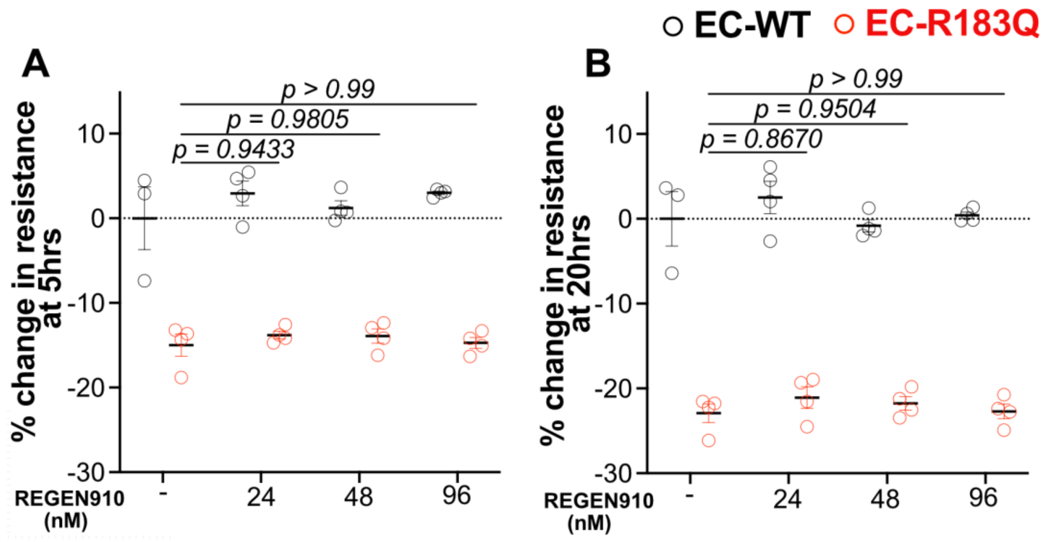


*Supplemental Figure 9 – Regeneron (REGEN910) anti-ANGPT2 antibody had no effect on EC-R183Q barrier function.* EC-WT (black) and EC-R183Q (red) were plated in TEER plates and treated with different concentrations of anti-ANGPT2 antibody (24nM to 96nM). Quantification of percent change in endothelial barrier (**A**) at 5 hours (**B**) at 20 hours. n= 4 electrodes/condition were measured. 3 independent experiments were performed. The p-values were calculated by Brown-Forsythe and Welch ANOVA test followed by Dunnett T3 multiple comparison test.

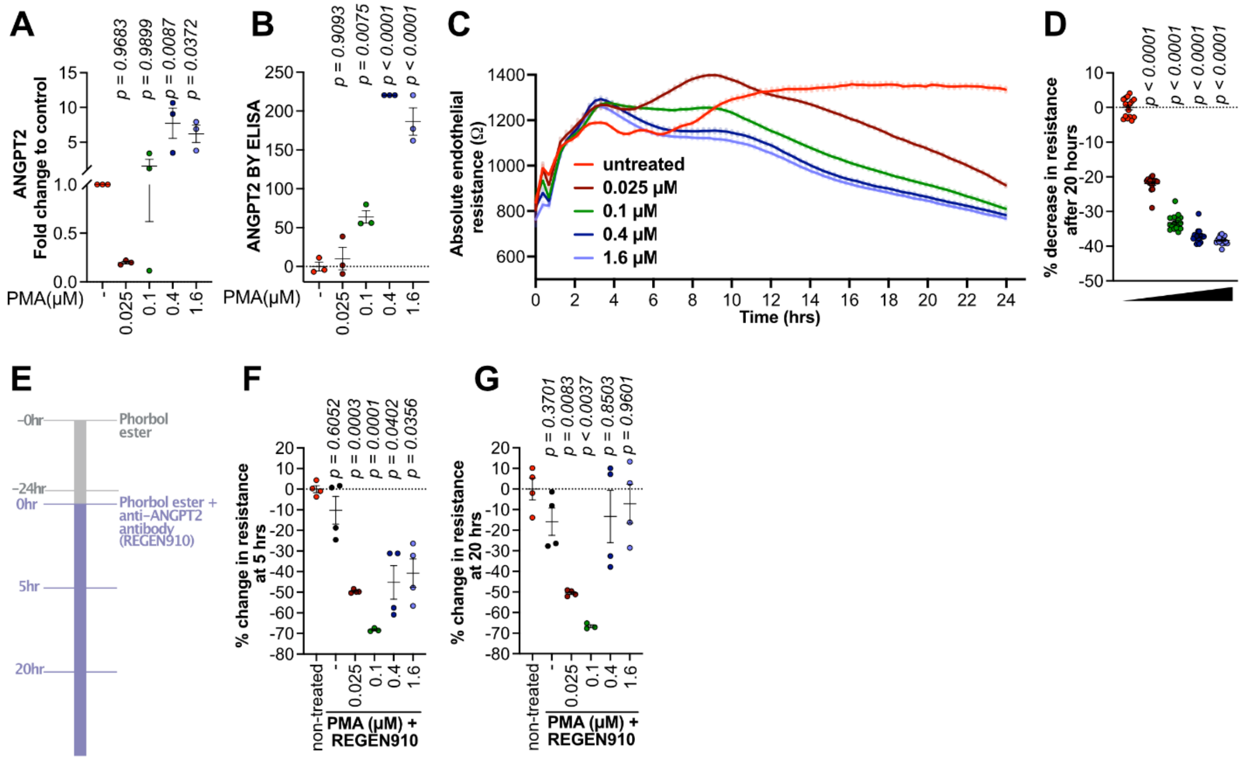


*Supplemental Figure 10 – Verification of REGEN910 activity.* Human endothelial colony forming cells (ECFC) were incubated with increasing concentrations of Phorbol 12-myristate 13-acetate (PMA) for 16 hours. ANGPT2 measured by (**A**) qPCR or by (**B**) ELISA increased with increasing concentrations of PMA. (**C**) Absolute endothelial resistance was measured by TEER in ECFC treated with increasing concentrations of PMA. (**D**) shows the percentage change in resistance at 20 hours at each dose of PMA. (**E**) Schematic of TEER assay with PMA (0.025-1.6μM) and anti-ANGPT2 (REGEN910) at 16nM. Quantification of percentage change in resistance at (**F**) 5 hours and (**G**) 20 hours. REGEN910 restored TEER in PMA treated cells indicating it was effective in blocking ANGPT2. n= 4. 3 independent experiments were performed. The p-values were calculated by Brown-Forsythe and Welch ANOVA test followed by Dunnett T3 multiple comparison test.

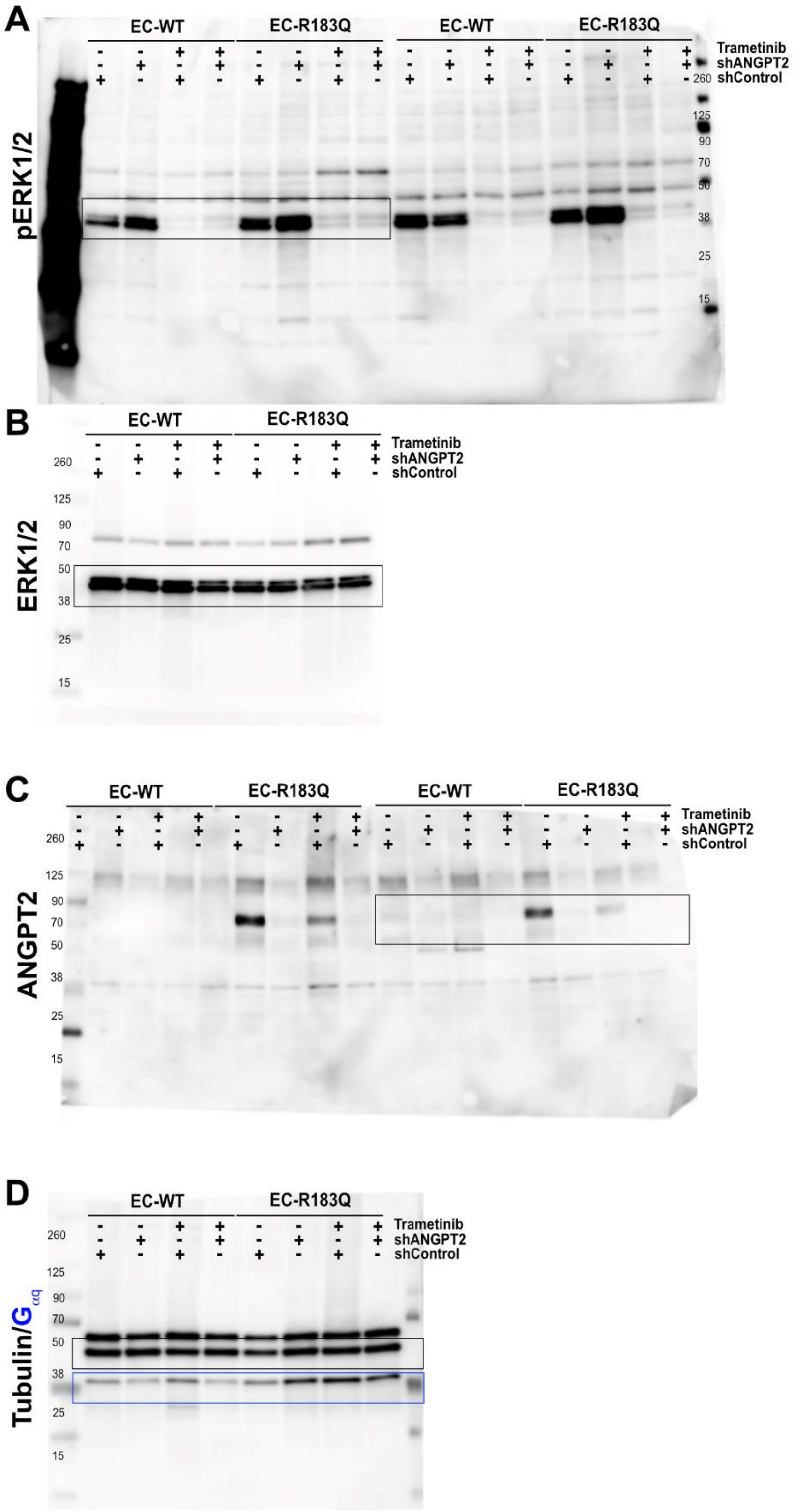


*Supplemental Figure 11 - Uncut WBs from combination of shANGPT2 and trametinib.* WBs probed with (**A**) anti-pERK1/2 (**B**) anti-ERK (**C**) anti-ANGPT2 (**D**) anti-tubulin and anti-$G_{\alpha q}$. The catalog numbers and details for each antibody are provided in the *Major Resource Table*. *Table*. Molecular weight markers are shown on the left side of each membrane.

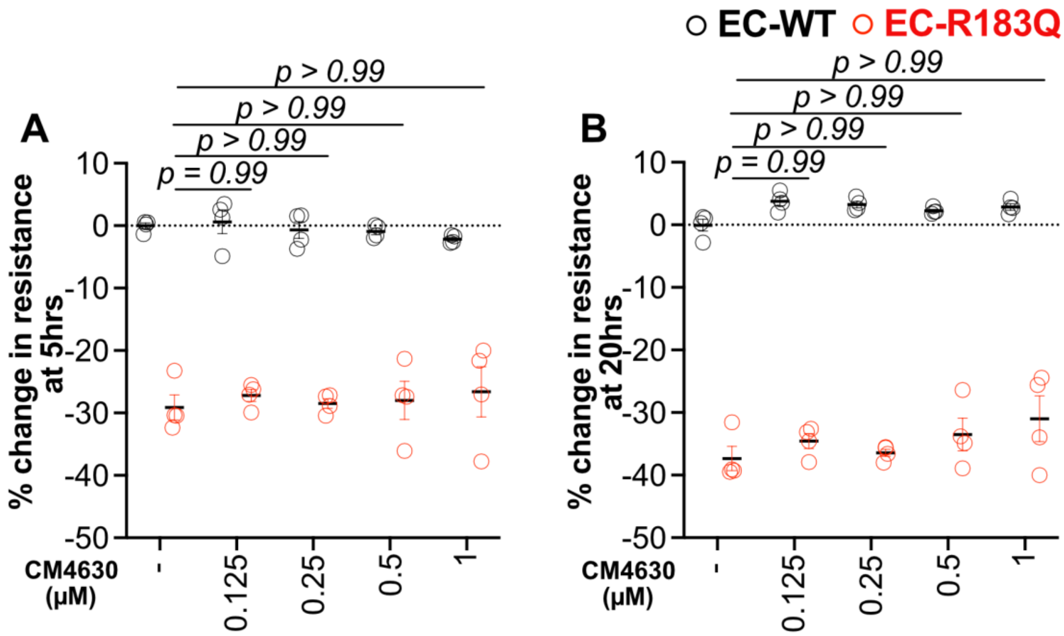


*Supplemental Figure 12 – CRAC channel inhibition using CM4620 has no effect on mutant GNAQ R183Q barrier function.* EC-WT (black) and EC-R183Q (red) were plated with different concentrations of CM4620 (0 to 1μM) in TEER plates and absolute endothelial resistance (Ω) was measured at 4000Hz. Quantification of percent change in endothelial barrier (**A**) at 5 hours (**B**) at 20 hours. n= 4. 3 independent experiments were performed. The p-values were calculated by Brown-Forsythe and Welch ANOVA test followed by Dunnett T3 multiple comparison test.

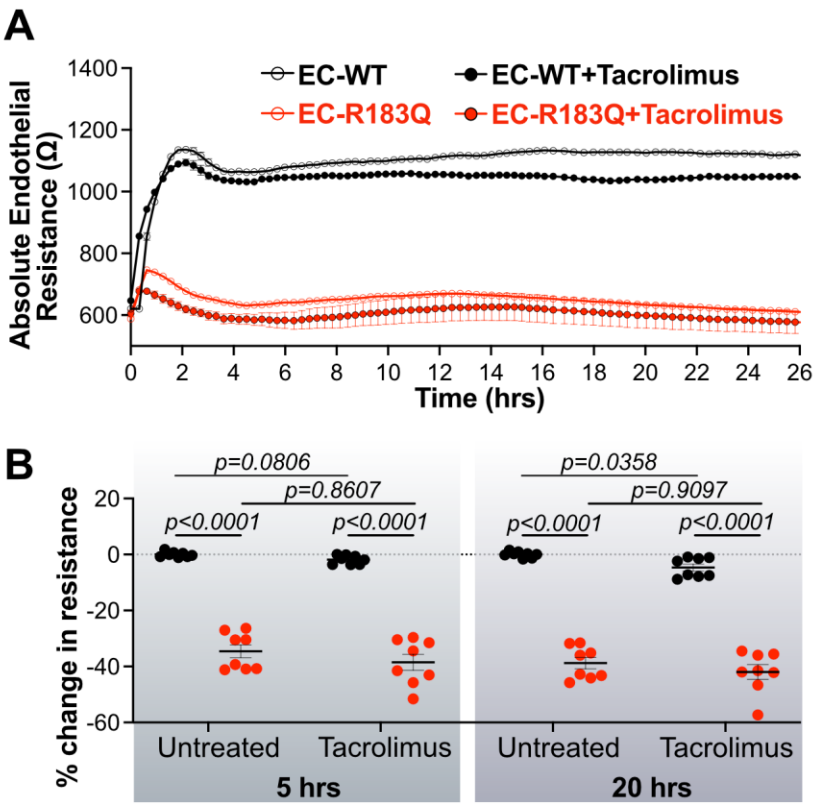


*Supplemental Figure 13 – Calcineurin inhibitor, Tacrolimus, had no effect on mutant GNAQ R183Q barrier function.* (**A**) Absolute endothelial resistance (Ω) of EC-WT (black open circles), EC-R183Q (red open circles), EC-WT treated with Tacrolimus (black closed circles), EC-R183Q treated with Tacrolimus (red closed circles) over 24 hours. (**B**) Quantification of percent change in resistance at 5 hours and 20 hours EC-WT and EC-R183Q ± Tacrolimus (200nM). n= 8. 3 independent experiments were performed. The p-values were calculated by Brown-Forsythe and Welch ANOVA test followed by Dunnett T3 multiple comparison test.

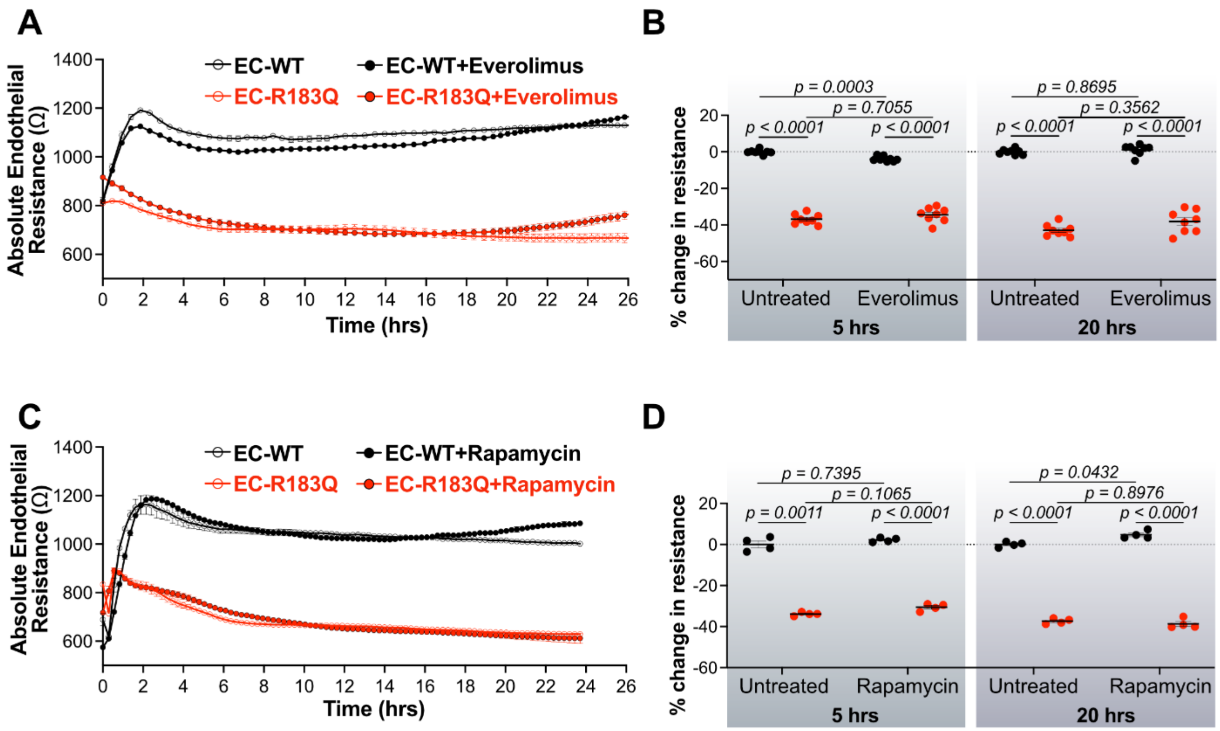


*Supplemental Figure 14 – mTOR inhibitors, Everolimus and Rapamycin, have no effect on mutant GNAQ R183Q barrier function.* (**A**) Absolute endothelial resistance (Ω) of EC-WT (black open circles), EC-R183Q (red open circles), EC-WT treated with Everolimus (black closed circles), EC-R183Q treated with Everolimus (red closed circles) over 24 hours. (**B**) Quantification of percent change in resistance at 5 hours and 20 hours EC-WT and EC-R183Q ± Everolimus (200nM). n= 8. (**C**) Absolute endothelial resistance (Ω) of EC-WT (black open circles), EC-R183Q (red open circles), EC-WT treated with Rapamycin (black closed circles), EC-R183Q treated with Rapamycin (red closed circles) over 24 hours. (**D**) Quantification of percent change in resistance at 5 hours and 20 hours EC-WT and EC-R183Q ± Rapamycin (20nM). n= 4. 3 independent experiments were performed. The p-values were calculated by Brown-Forsythe and Welch ANOVA test followed by Dunnett T3 multiple comparison test.
